# Climate and management changes drove more stress-tolerant and less competitive plant communities in 40 years

**DOI:** 10.1101/2023.08.23.554452

**Authors:** Marie-Charlotte Bopp, Elena Kazakou, Aurélie Metay, Jacques Maillet, Marie-Claude Quidoz, Léa Genty, Guillaume Fried

## Abstract

Spontaneous plant communities have undergone considerable constraints due to human-mediated changes. Understanding how plant communities are shifting in response to land management and climate changes is necessary to predict future ecosystem functioning and improve the resilience of managed ecosystems, such as agroecosystems. Using Mediterranean weed communities as models of managed plant communities in a climate change hotspot, we quantified to which extent they have shifted from the 1980s to the 2020s in response to climate and management changes in vineyards. In four decades, the annual range of temperatures (*i.e.* the difference between the warmest month’s and the coldest month’s mean temperatures) increased by 1.2°C and the summer temperatures by 2°C. Weed management diversified over time with the adoption of mowing that replaced the chemical weeding of inter-rows. Current weed communities were 41% more abundant, 24% more diverse and with a less even distribution of abundance across species than the 1980s communities at the vineyard level. Modern communities were composed of more annual species (57% of annual species in the 1980s versus 80% in the 2020s) with lower lateral spreadability and seed mass and were composed of fewer C4 species. They had higher community-weighted specific leaf area, higher leaf dry matter content and lower leaf area than 1980s weed communities. At the community level, the onset of flowering was earlier and the duration of flowering was longer in the 2020s. Climate change induced more stress-tolerant communities in the 2020s while the diversification of weed management practices over time filtered less competitive communities. This study shows that plant communities are adapting to climate change and that land management is a strong lever for action to model more diverse and functional plant communities in the future.

## INTRODUCTION

Spontaneous plant communities have undergone considerable constraints due to human-mediated changes at global (*e.g.* climate change) and regional scales (*e.g.* land use change) (Inderjit et al. 2017). At the global scale, climate change results in higher temperatures, altered precipitation, higher frequency of extreme weather events and increased level of CO_2_ (Peters, Breitsameter, and Gerowitt 2014). The Mediterranean area has been identified as a ‘climate change hotspot’, exceeding global warming average rates by 20% and having reduced rainfall, especially in summer (Ali et al. 2022). Consequently, climate change is expected to impact disproportionally Mediterranean vegetation compared to other regions of the world (Newbold et al. 2020). The projections of the effect of climate change on plant communities are severe: decreased species richness, reduction of the species range, shifts toward a higher proportion of annual species in communities and a lower proportion of forbs (Pfeifer-Meister et al. 2016; Sala et al. 2000). These changes in plant composition can ultimately alter many ecosystem processes (Cardinale et al. 2012). This changing global filter also interacts with regional filters like human management (Bourgeois et al. 2021) which have become more pronounced during the last decades. In the last 50 years, agriculture intensification including herbicides and mineral fertilizers uses has exerted strong selection pressures on spontaneous plant communities, favouring communities adapted to a high level of disturbance and resources (Bourgeois et al. 2019). As a result of this more intensive weed management, the diversity of weed communities has also dramatically decreased in arable lands (Fried, Chauvel, and Reboud 2009; Cirujeda, Aibar, and Zaragoza 2011; Fried, Dessaint, and Reboud 2016; Storkey et al. 2011; 2021).

In agroecosystems, the temporal changes in plant communities were mostly described in annual cropping systems while few studies have investigated spontaneous flora shifts in less disturbed systems like perennial cropping systems. We assume that Mediterranean vineyard weed communities are particularly relevant models to study flora shifts because the agricultural practices of vineyards strongly evolved from the late 1970s to nowadays. At the end of the 1970s, herbicides started to be widely applied on both the rows and the inter-rows of vines (*i.e.* the free space between the rows of vines) (Maillet 1981). This constituted an abrupt change in environmental conditions resulting in drastic changes in weed communities (Maillet 1992). In the 2000s, the awareness of the detrimental effects of herbicides on the environment (*e.g.* water quality) forced farmers to limit their use. Superficial tillage and mowing gradually replaced herbicide use in the inter-rows, leading to more integrated weed management (Fernández-Mena et al. 2021). In addition, regulations are increasingly restricting the use of herbicides, with, for example, a ban on the use of glyphosate in the inter-rows of French vineyards from 2021 (ANSES 2020). However, chemical weeding of the rows is still possible and frequent in French Mediterranean vineyards (Bopp, Fried, et al. 2022).

Trait-based approaches provide a relevant framework to better understand how weed communities shift in response to climate and management changes (Richner et al. 2015). Functional trait values at the community level, also known as ‘functional markers’ (Garnier et al. 2004), respond to environmental filtering (*e.g.* climate change) (Lavorel and Garnier 2002) and anthropogenic drivers (*e.g.* agricultural practices) (Booth and Swanton 2002; Navas 2012; Damour, Navas, and Garnier 2018). This approach assumes that only certain species with specific trait values will persist within communities in particular climate and management conditions. Synchronic Mediterranean studies of trait patterns along aridity gradients in natural environments suggested that climate change filter communities with a more resource-conservative strategy: communities with small species, low leaf area and specific leaf area (SLA) and populations with short flowering duration (Costa-Saura et al. 2016; Nunes et al. 2017; de la Riva et al. 2018). Moreover, high temperatures and frequent drought would favour C4 weed species (Korres et al. 2016), late-emerging and thermophilic species (Peters, Breitsameter, and Gerowitt 2014) while rising CO2 atmospheric pressure is expected to favour C3 species. Thus, the impact of climate change on photosynthetic pathways is not entirely clear (Peters, Breitsameter, and Gerowitt 2014). Yet, few studies have revealed how climate change might modify plant communities using diachronic approaches (Feeley et al. 2020; Harrison, Spasojevic, and Li 2020).

Weed management practices can be considered as anthropogenic disturbances and can be classified according to their intensity and frequency (Kazakou et al. 2016; Gaba et al. 2014). For instance, chemical weeding and tillage represent both high-intensity disturbances as they destroyed the whole aboveground biomass (Kazakou et al. 2016). In contrast, mowing might be considered a lower-intensity disturbance as this practice removes the aboveground biomass partially (MacLaren, Bennett, and Dehnen-Schmutz 2019). Recent synchronic studies (*i.e.* comparing simultaneously vineyards with different managements) demonstrated the strong filtering effects of management on weed communities in vineyards (Kazakou et al. 2016; MacLaren, Bennett, and Dehnen-Schmutz 2019). More precisely, mowing favoured more weed communities composed of conservative and competitive species, *i.e.* with more perennials, higher seed mass, higher stature, lower specific leaf area (SLA) and higher leaf dry matter content (LDMC) than communities managed with other practices like tillage (Hall et al. 2020; Mainardis et al. 2020; Kazakou et al. 2016; Guerra et al. 2021; Bopp, Kazakou, et al. 2022; Fried et al. 2022). In contrast, tillage and chemical weeding favoured communities composed of acquisitive species, *i.e.* annual species with high SLA and low LDMC and late-flowering populations (Hall et al. 2020; Bopp, Kazakou, et al. 2022; Fried et al. 2022). However, herbicides were shown to favour contrasted life forms in weed communities: both annual and some perennial species (*e.g.* geophytes) (Hall et al., 2020). The CSR scheme of Grime (1977) is an integrative approach to studying adaptative strategies that species use to cope with two constraints: (i) stress related to lower resource availability (*e.g.* water stress related to climate change in the Mediterranean area) and (ii) disturbance related to the sudden destruction of biomass (*e.g.* weed management practice as tillage, mowing, chemical weeding). The CSR scheme describes three main species strategies: (i) stress tolerators (S) in resource-poor habitats with low disturbance, (ii) ruderals (R) in resource-rich and highly disturbed environments (*e.g.* agroecosystems) and (iii) competitors (C) in highly productive habitats with low-stress intensity and disturbance. If weed species are mostly ruderals as cultivated habitats are by definition regularly disturbed by tillage (Mahaut et al. 2020; Bourgeois et al. 2019), a recent study demonstrated that the proportion of CSR strategies in weed communities changed according to the weed management used (Fried et al. 2022). For instance, mowing favoured communities with more competitive strategies while tillage favoured species with more ruderal strategies. Climate change might also modify the relative proportion of species’s CSR strategies within the communities over time (MacLaren et al. 2020). For instance, under climate change, weed species with stress tolerator strategies are more likely to occur (Korres et al. 2016). If recent synchronic studies exist in weed functional response to management (*e.g.* MacLaren et al. 2019, Hall et al. 2020) or to climate (Bopp, Kazakou, et al. 2022), no study, to our knowledge, has yet been conducted on weed community functional changes over time in vineyards.

In this study, we quantified how climate and management changes have shifted weed communities from the 1980s to the 2020s, using a historic network of 40 Mediterranean vineyards in South France. First, we assessed climate change intensity and characterise changes in weed management practices in the vineyard network. Then, we described shifts in taxonomic and functional weed community structures after four decades using 374 floristic surveys that were performed during two years at each period (1978, 1979, 2020, 2021) and at 3 seasons (March, June, October). Finally, we tested if climate conditions and weed management practices drove changes in weed community structure between the 1980s and the 2020s. We hypothesised that climate change would favour species with more water-conservative traits (*e.g.* C4 metabolism, small leaf area, high LDMC) and stress-tolerant strategies. Moreover, we expected that the decrease in herbicide use replaced by mowing would favour more conservative and competitive strategies.

## MATERIAL AND METHODS

### 1. Climate, soil and weed management characterisation of the historical vineyard network

The network was composed of 40 vineyards located around Montpellier, France (Figure 1). This network was first set up in 1978 to study weed management effects on weed community composition (Maillet 1981). The vineyards are located in a Mediterranean climate characterised by warm temperatures (14°C mean annual temperatures on average from 1978 to 2021), irregular rainfall (814 mm per year) and dry summers (88% of annual precipitation occurs during the other seasons). Soil textures of vineyards varied along a North-East/South-West gradient determined by the distance to the Mediterranean Sea: from sandy vineyards nearby the Sea to loamy and clay/loamy soils in further northeast vineyards.

**Figure 1.**
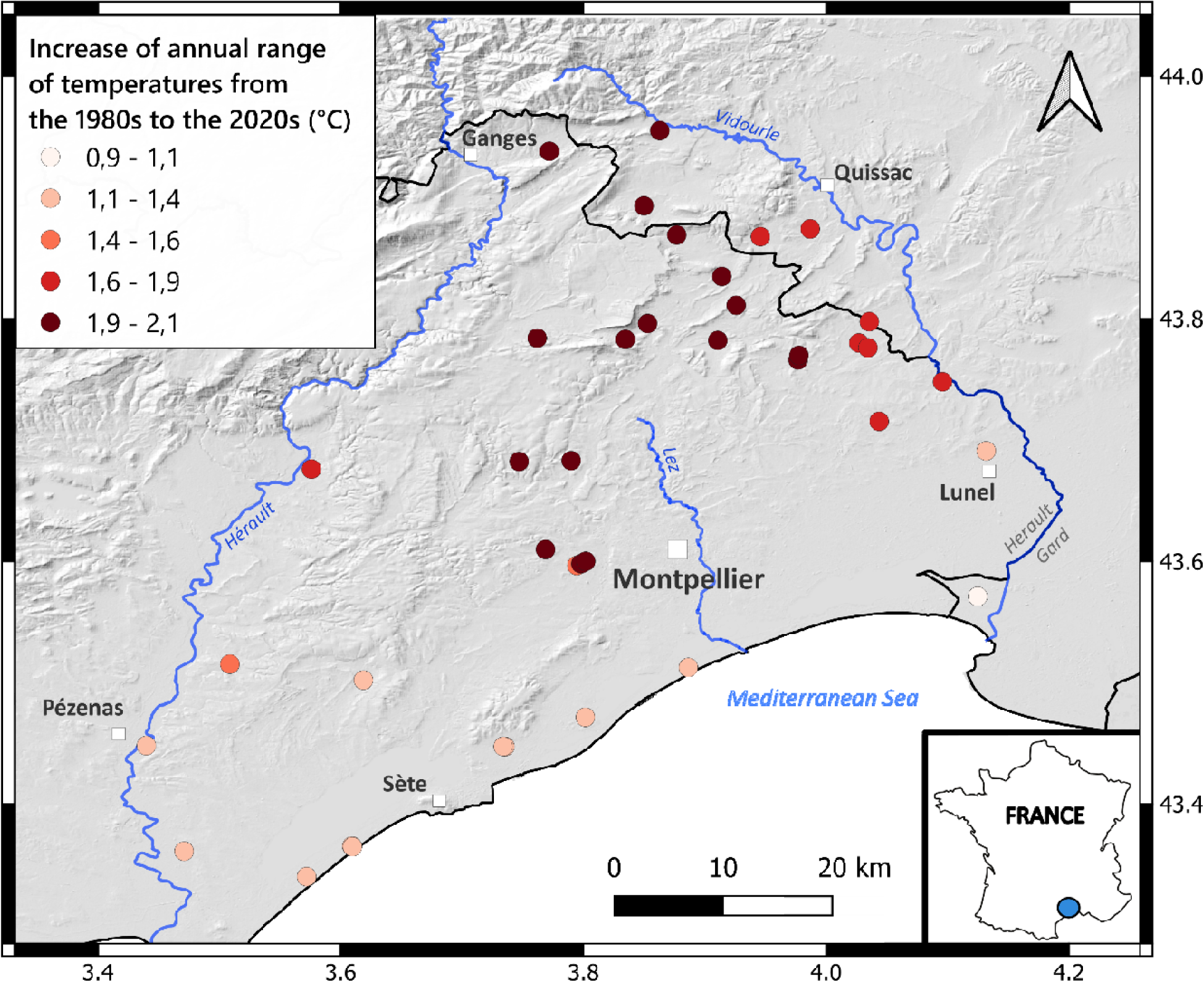
Geographical distribution of the increase of annual range of temperatures from the 1980s to the 2020s experienced by the 40 vineyards of the network, Montpellier, France. Each dot is a vineyard and its colour represents the extent of the increase in the annual range of temperatures (from light red for low increases to dark red for high increases). The dark lines represent the department’s outlines. The blue lines represent the main rivers of the region. The values displayed on the frame surrounding the map are the latitudinal and longitudinal coordinates.

To characterise climate change, variables describing seasonal and annual climate were extracted from the estimations obtained by the SAFRAN interpolation method (Analysis system providing atmospheric information to the snow) on an 8 km × 8 km grid (Durand et al. 1993) from 1971 to 2021. Eleven variables were selected: mean seasonal temperatures and rainfall (spring, summer, autumn and winter), growing season length (GSL) (*i.e.* the number of days in a year with daily mean temperature above 5°C during at least 6 consecutive days), numbers of frost days and annual range of temperatures (*i.e.* the difference between the warmest month’s and the coldest month’s mean temperatures), and were averaged over 5 different decades from 1971 to 2021 (1971-1980; 1981-1990; 1991-2000; 2001-2010; 2011-2021). Due to the SAFRAN grid, 31 neighbouring vineyards on 40 shared the same climate data at least with another vineyard: 20 squares of 8x8 km² with different climate data were used for the 40 vineyards of the network.

To characterise changes in weed management practices of the rows and the inter-rows from 1978 to 2021, we conducted two interviews in 2020 and 2021 with each farmer of the network (see Appendix 3: Section S1). We considered the five following binary practice variables: (i) chemical weeding of rows, (ii) chemical weeding of inter-rows, (iii) tillage of rows, (iv) tillage of inter-rows and (v) mowing of inter-row. Each variable had two possible values: 0, not applied; 1, applied. To define the trajectory of the weed management practices of each vineyard, we used the values of these five variables over five different periods: 1978-1979 (the initial weed management practices); 1980-1990; 1991-2000; 2001-2010 and 2011-2021 (the final weed management practices).

### 2. Weed community composition in the 1980s and the 2020s

Weed community composition was determined in 1978 and 1979 (as ‘1980s communities’ hereafter) (Maillet 1981) and in 2020 and 2021 (as ‘2020s communities’) at 3 different seasons (late winter, early summer and autumn) (Appendix S1: Figure S1). In total, the dataset was composed of 374 floristic surveys (for more details, see Appendix S1: Figure S1). To compare the floristic compositions over the two periods, we applied the same method of floristic surveys. In each vineyard, a rectangular area of 400 m² (40m x 10m) was delimited (4 inter-rows and 3 rows of the same length). To estimate species abundance, we used five abundance classes developed in Barralis (1976): ‘1’, less than 1 individual/m²; ‘2’, 1–2 individuals/m²; ‘3’, 3–20 individuals/m²; ‘4’, 21–50 individuals/m²; ‘5’, more than 50 individuals/m². We transformed these scores into a quantitative scaling using the median of the range of each density class as followed: ‘1’, 0.5 individual/m²; ‘2’, 1.5 individuals/m²; ‘3’, 11.5 individuals/m²; ‘4’, 35.5 individuals/m²; ‘5’, 75 individuals/m². A list of species and distinct abundance scores were noted at the plot level. To control the ‘observer effect’, we conducted several floristic surveys in 2020 with Jacques Maillet, who conducted the floristic surveys in the 1980s. No major biases were observed during these floristic surveys. For each community, we computed the richness, the abundance (*i.e.* the sum of the density of each species), the Shannon diversity index and Pielou’s evenness index which is the ratio between the Shannon index and the logarithm of the community richness (Pielou 1966).

### 3. Functional properties of the 1980s and 2020s weed communities

Seven plant traits, two phenological population-level traits and four Ellenberg indices were selected to capture plant responses to climate change and weed management changes. Three traits of the Leaf-Height-Seed (LHS) strategy scheme were selected (Westoby, 1998): 1) specific leaf area (SLA, m².kg^-1^) related to the speed of resources acquisition (Wright et al., 2004), 2) maximum height (m) related to light and nutrient acquisition (Westoby et al., 2002), 3) seed mass (g) which represents the “colonisation-competition” trade-off (Moles and Westoby, 2006). Leaf area (m²) was also selected as a good *proxy* for the competitive abilities of plants and lateral spread as a response trait to disturbance. This qualitative trait represents species’ abilities to develop horizontally (species with rhizomes or forming tussocks); it is rated as followed: “1”, therophytes; “2”, perennials with compact unbranched rhizomes or forming small tussocks (less than 100 mm in diameter); “3”, perennials with a rhizomatous system or tussocks reaching from 100 to 250 mm; “4”, perennials reaching a diameter of 251 to 1000 mm. Leaf Dry Matter Content (*i.e.* the ratio between leaf dry mass and leaf fresh mass) was selected as a *proxy* for plant investment in leaf structure. Photosynthesis pathways (C3/C4) were assessed to identify the species with water-saving photosynthesis (Pyankov et al. 2010). Phenology was characterised by the months of flowering onset and the number of months of flowering determined at the population level for each species. Species resource requirements were characterised through four Ellenberg indices: light (EIV_L), temperature (EIV_T), soil moisture (EIV_F) and nitrogen (EIV_N). Raunkiær plant life forms were included.

Leaf area, SLA and LDMC were measured on the weed species present in the 2020s communities (8 individuals per species). The other traits were extracted from different databases: Seed Information Database for seed mass (Royal Botanic Gardens Kew 2021), *Flora Gallica* for maximum height (Tison and De Foucault 2014), lateral spread from Hodgson et al. (1995) supplemented by expert opinion (G. Fried, pers. com.), Baseflor for flowering onset, duration of flowering and the four Ellenberg indices (Julve 1998). Flowering onset and duration of flowering are quantified at the population level and not at the individual level (*i.e.,* a long flowering duration does not mean that individuals of that species can flower during a long period but that different individuals of that species cover together a long period of flowering). Photosynthetic pathways were extracted from Pyankov et al. (2010).

To scale up the trait values from the species level to the community level, we calculated Community Weighted Means (CWM, *i.e.* the average of the trait values of species present in a community, weighted by their relative abundances) (Garnier et al. 2004) using at least 80% of the most abundant species of the weed communities in the calculation (Mass ratio hypothesis, (Grime 1998)). We calculated median values for lateral spread and the percentage of C4 species’ abundance within the communities. Based on the CWM of SLA, LDMC and leaf area, we calculated the competitiveness (C), stress-tolerance (S) and ruderal (R) scores following Pierce et al. (2017) and Li and Shipley (2017).

### 4. Data analyses

#### a) Description of climate, weed management and weed community functional structure using multivariate analyses

Multivariate analyses were performed to characterise climate and weed management changes per decade from the 1980s to the 2020s (1980s, 1990s, 2000s, 2010s and 2020s) (see Appendix S3: Section S2). Four categories of the 2020s weed management were defined using a Principal Component Analysis (PCA) and hierarchical clustering (see Appendix S1: Figure S2, Appendix S3: Section S3). Cluster ‘Chem’ gathers four vineyards with chemical weeding of both rows and inter-rows. Cluster ‘Till.IR.Chem.R’ regroups 19 vineyards with tilled inter-rows and chemically weeded rows. Cluster ‘Mow.IR.Till.R’ gathers 11 vineyards with mowed inter-rows and tilled rows. Cluster ‘Till’ groups six vineyards with tilled rows and inter-rows. A PCA was also performed on the CWM of the 13 traits of the 1980s and 2020s communities. To select a parsimonious number of dimensions which represents well the functional space, we used the elbow inflexion point method on the Area Under the Curve (AUC) and the Mean Absolute Deviation (MAD) following Mouillot et al. (2021).

#### b) Comparison of structure and composition of the 1980s and 2020s weed communities

Two different datasets were used for data analyses (Appendix S1: Figure S1). The first one is the ‘full dataset’ composed the 374 floristic surveys from the 1980s and the 2020s. This dataset was used to test the significance of the difference in means of taxonomic indices (abundance, richness, Shannon, Pielou), Raunkiær plant life forms, CWM and CSR scores over time. Using the full dataset (n=374), we fitted linear mixed models with the period (the 1980s/2020s) as a fixed effect and the season (March/June/October), the year (1978, 1979, 2020, 2021) and plot identity as random effects. An ANOVA was performed on these models to test fixed effect significance. Post-hoc Tukey and Wilcoxon tests were then computed to assess the direction of the changes in functional properties of weed communities from the 1980s to the 2020s. Moreover, the changes in abundance and frequency of species over time were quantified using indices of relative changes following this formula: (Indice_2020s_ – Indice_1980s_) / (Indice_2020s_ + Indice_1980s_). From these indices, we identified (i) the species that increased in abundance and frequency defined as the species with more than 0.75 of relative changes in abundance and frequency, (ii) the species that decreased in abundance and frequency with less than -0.75 of relative changes and (iii) the species that remained stable in abundance and frequency with relative changes within the range of [-0.75; 0.75].

#### c) Identification of the drivers of weed community structure shifts over time

The second dataset was built to test whether climate and management changes explained the changes in taxonomic and functional properties of weed communities (Appendix S1: Figure S1). To build an ‘explaining’ dataset, we averaged the functional and taxonomic metrics over the two years for each period: from 5 floristic surveys per period (*e.g.* for the 1980s, March 1978, 1979, June 1978, 1979 and October 1979) to 3 floristic surveys per period (*e.g.* March 1980s, June 1980s, October 1980s). Then, each season of each period was comparable with the season of the other period (*e.g.* March 1980s with March 2020s). This ‘explaining’ dataset was composed of 120 taxonomic and functional shifts of communities from the 1980s to the 2020s: 40 vineyard community shifts x 3 seasons (Appendix S1: Figure S1). After removing missing data, the dataset consisted of 108 functional shifts. We used two different ways of computing indexes of changes: (i) for comparing coordinates of the CWM in the functional PCA, we computed the difference between the coordinates in the 2020s and the coordinates in the 1980s for each selected axes of the multivariate analysis (Appendix S2: Table S1) and (ii) for comparing ‘raw values’ of different metrics (CWM, CSR scores, annual temperature range, abundance, richness, Shannon and Pielou indices), we computed indices of relative changes using the following formula: (Indice_2020s_ – Indice_1980s_) /(Indice_2020s_ + Indice_1980s_) (Appendix S2: Table S1).

We selected the climate and management drivers that explained most of the climate and management changes over the four decades from the multivariate analyses based on climate and management characteristics per decade. As a *proxy* for climate change, we selected the relative changes in the annual T°C range between the 1980s and the 2020s (*i.e.* the difference between the warmest months and the coldest month’s mean temperatures). To describe weed management trajectory, we used three variables: (i) the number of decades since herbicides were stopped, (ii) the number of decades since mowing was used and (iii) the additional number of different management practices applied at the plot level from the 1980s to the 2020s, as a *proxy* for the trajectory toward more integrated weed management (*e.g.* if tillage was the only management practice used in the 1980s and if mowing and tillage were used in 2020s, the value would be ‘1’ additional weed management practice applied). To describe current weed management, we selected the categorical variable describing 2020s weed management practices with four different groups (**‘Chem’**: vineyards with chemical weeding of both rows and inter-rows; **‘Till.IR.Chem.R’**: vineyards with tilled inter-rows and chemically weeded rows; **‘Mow.IR.Till.R’**: vineyards with mowed inter-rows and tilled rows; **‘Till’:** vineyards with tilled rows and inter-rows) (Appendix S2: Table S1). Using the ‘explaining’ dataset (n=108), we fitted linear mixed models to explain the shifts in CWM, taxonomic indices (abundance, richness, Pielou, Shannon) and CSR scores over time by the selected drivers of climate and management changes as fixed effects and with the season and plot identity as random effects. The selection of models was done based on the corrected Akaike Information Criterion (AICc) with the *dredge* function from the *MuMIn* R package (Burnham and Anderson 2002). All data analyses were carried out using R version 4.1.1 (R Core Team 2021).

## RESULTS

### 1. Climate and management changes from the 1980s to the 2020s

#### a) Climate change from the 1980s to the 2020s nearby Montpellier, France: the annual range of temperature increased by 1.2°C

Over the two periods, seasonal climate variables (spring, summer, autumn, winter temperature, spring, summer, autumn, winter rainfall, and the annual number of frost days) of the vineyards were well explained by the first two PCA axes (84.7% of total variance) (Figure 2). The first axis explained most of the climate variance (60.5%) and intra-network variations across the vineyards from the same decades. This axis opposed vineyards located in warmer and drier areas with longer growing season length, located near the coastline to vineyards located in areas with higher rainfall, higher numbers of frost days and shorter growing season length, located in the northeast of the network. This axis also represented a slight decrease in seasonal precipitations over the decades (Appendix S1: Figure S3). The second axis described mostly inter-decades variations of the climate conditions (24.2% of explained variance) (Figure 2, Appendix S1: Figure S3). This axis was mainly driven by the annual range of temperatures (*i.e.* the difference between the mean temperatures of the warmest month and the coldest month), the summer mean temperature and the number of frost days. In 40 years, the annual range of temperatures increased by 1.2°C (from 13.5°C averaged over the 1971-1980 decade to 14.7°C averaged over the 2011-2021 decade). Moreover, summer mean temperatures have increased by 2°C in 40 years (21°C in the 1980s and 23°C in the 2020s) and spring temperatures by 1.4°C (12°C in the 1980s to 13.4°C in the 2020s) (Table 1). The annual number of frost days doubled in 40 years: from 11 days in the 1980s to 23 in the 2020s (Table 1). Within the vineyard network, the increase in the annual range of temperatures was heterogeneous and varied from an increase of 0.9°C (vineyards located nearby the Mediterranean Sea) to an increase of 2.1°C (Northern vineyards) (Figure 2).

**Figure 2.**
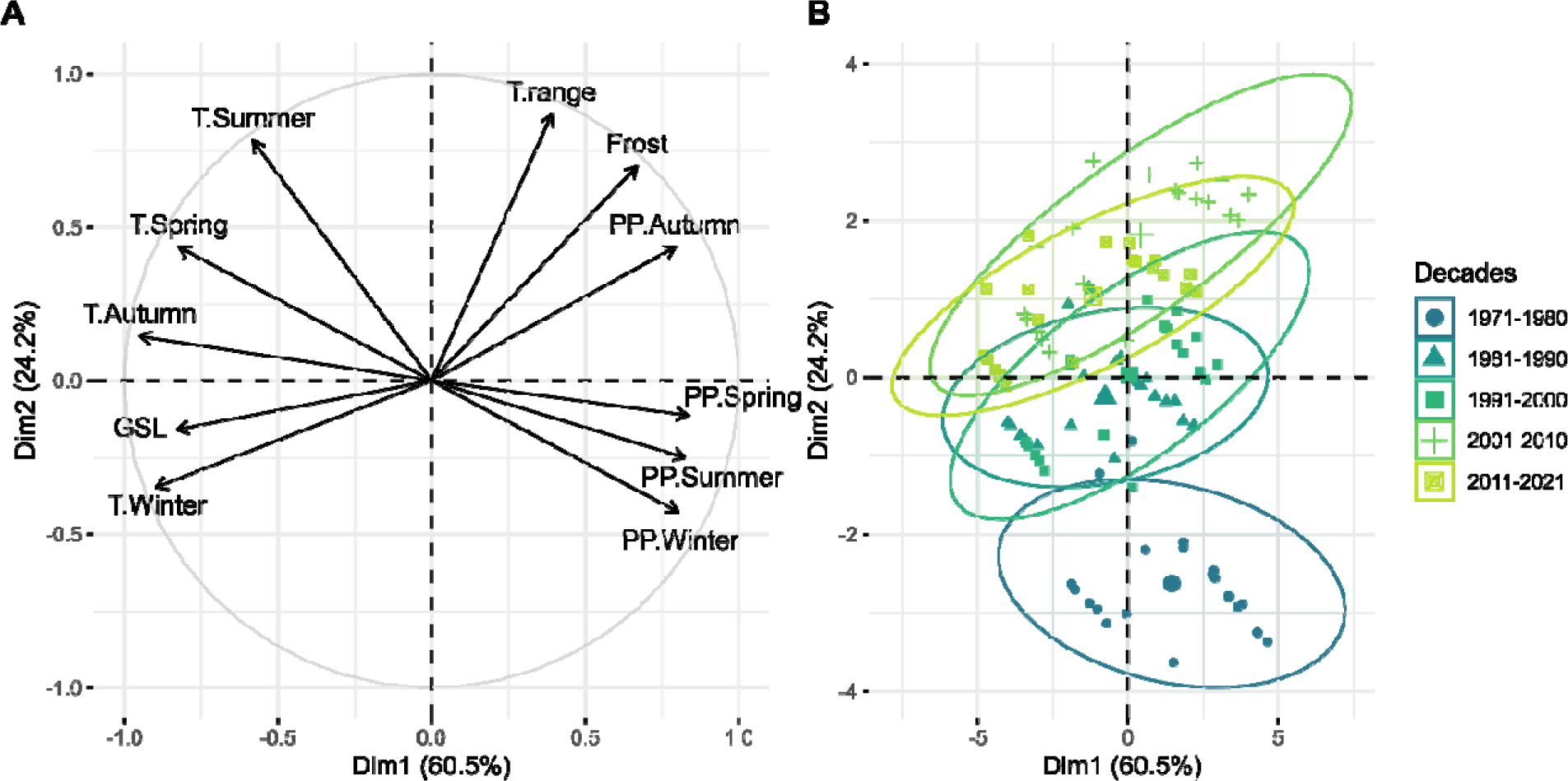
Climate change from the 1970s to the 2020s. **A)** Correlation circle of the climate variables on the first two axes of the climate PCA over the decades from the 1970s to the 2020s. **B)** Vineyard projection on the first two axes of the climate PCA for each decade from the 1970s to the 2020s. Each variable was averaged over the five studied decades: 1971-1980, 1981-1990, 1991-2000, 2001-2010, 2011-2021. T.Summer, mean summer temperature (°C); T.Spring, mean spring temperature (°C); T.Autumn, mean autumn temperature (°C); T.Winter, mean winter temperature (°C); PP.Spring, winter cumulated rainfall (mm); PP.Autumn, autumn cumulated rainfall (mm); PP.Winter, winter cumulated rainfall (mm); PP.Summer, summer cumulated rainfall (mm); T.range, annual temperature range (°C); Frost, number of frost days; GSL, growing season length.

**Table 1.** Averaged climate and weed management characteristics in the 1980s and the 2020s. Minimum and maximum values are reported in brackets.

| **Drivers** | **Variables** | **Unit** | **1980s** | **2020s** |
| --- | --- | --- | --- | --- |
| **Climate** | Annual temperature range | °C | 15.7 (15-16.9) | 17.4 (16.2-18.1) |
|  | Growing season length | Days | 339 (328-353) | 354 (337-363) |
|  | Number of frost days | Days | 11 (5-16) | 23 (6-38) |
|  | Winter rainfall | mm | 312 (232-373) | 144 (98-179) |
|  | Spring rainfall | mm | 228 (169-273) | 207 (128-254) |
|  | Summer rainfall | mm | 125 (84-151) | 104 (71-127) |
|  | Autumn rainfall | mm | 265 (166-337) | 293 (187-362) |
|  | Winter temperature | °C | 7.1 (5.9-8.2) | 7.4 (6-8.8) |
|  | Spring temperature | °C | 12.0 (10.8-12.9) | 13.4 (12.2-14.5) |
|  | Summer temperature | °C | 21.0 (19.9-22.1) | 23.1 (22.1-24.2) |
|  | Autumn temperature | °C | 13.8 (12.6-14.8) | 15.2 (13.8-16.5) |
| **Management** | Chemically weeded inter-rows | % of vineyards | 35 | 28 |
|  | Tilled inter-rows | % of vineyards | 63 | 95 |
|  | Mowed inter-rows | % of vineyards | 0 | 78 |
|  | Chemically weeded rows | % of vineyards | 70 | 68 |
|  | Tilled rows | % of vineyards | 23 | 60 |

#### b) Weed practice shifts in 40 years in Mediterranean vineyards: mowing adoption from the 2000s

The first two axes of the Principal Coordinates Analysis (PCoA) based on the weed management practice variables of decades from the 1980s to the 2020s explained 72.2% of the total variance (Figure 3). The first axis opposed weed management practices from chemical weeding of rows (rho: -0.69, *P* < 0.001) and inter-rows (rho: -0.77, *P* < 0.001) to tillage of rows (rho: 0.80, *P* < 0.001) and inter-rows (rho: 0.64, *P* < 0.001) and mowing of inter-rows (rho: 0.48, *P* < 0.001). The second axis discriminated the vineyards with low numbers of weed management practices used at the plot level (1 or 2) from the ones with more integrated weed management (*e.g.* 3 different weed management practices applied at the plot level) (Figure 3). More precisely, this axis opposed vineyards with tilled rows (rho: - 0.32, *P* < 0.001) and either chemical weeding or tillage of inter-rows to vineyards with a mixed combination of tilled (rho: 0.51, *P* < 0.001) and mowed inter-rows (rho: 0.69, *P* < 0.001) and chemical weeding of rows (rho: 0.66, *P* < 0.001). In the 1980s, farmers mostly tilled and applied herbicides. The mowing practice appeared in the 2000s, diversifying and increasing the number of weed management practices applied at the vineyard level. The median weed management practice that farmers applied in the 2020s was inter-row tilling and mowing and chemical weeding of rows (Figure 3). Thus, at the vineyard network level, the weed management shift from the 1980s to the 2020s was characterised by (i) less use of chemical weed control on the inter-rows and (ii) a combination of more weed management practices at the plot level due to the adoption of mowing from the 2000s (Figure 3).

**Figure 3.**
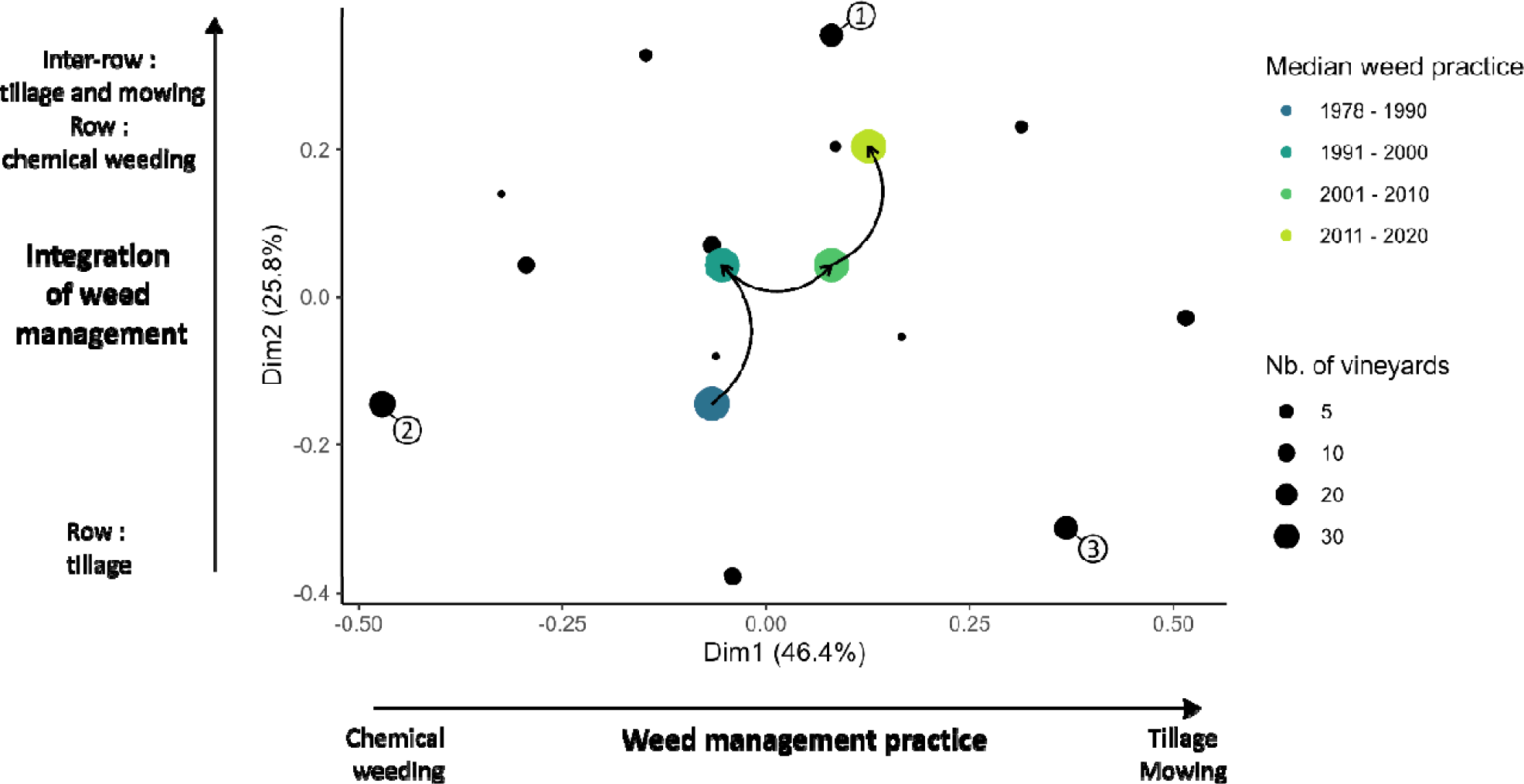
Changes in weed management practices from the 1980s to the 2020s in the Principal Coordinates Analysis (PCoA) space. Each dot represents a combination of weed practices at the plot level. The size of the black dots is proportional to the number of vineyards that share the same practice combination. The large coloured dots represent the median position of the vineyards of each decade in the PCoA space. The arrows illustrate the weed practice trajectory from the 1980s to the 2020s at the vineyard network scale. The first axis represents the weed management type used by the farmers. The second axis represents the number of weed practice types applied at the vineyard scale: from 1 practice type used (tillage of rows) to 3 practice types (tillage and mowing of inter-rows combined with chemically weeded rows). The vineyards located on position (1) in the practice PCoA are characterised by tillage and mowing of inter-rows and chemical weeding of the rows; the vineyards located on position (2) had chemically weeded rows and inter-rows; the vineyards located in position (3) have rows and inter-rows that are tilled. Nb. of vineyards, number of vineyards sharing the same weed management practices.

### 2. Shifts in weed community structure from the 1980s to the 2020s

In total, 436 species were found over the 374 floristic surveys of the two periods. At the vineyard network scale, a similar number of species were identified 40 years apart: 320 species in the 1980s and 319 species in the 2020s. At the vineyard scale, the mean species richness was higher in the 2020s (34 species/community) than in the 1980s (26 species/community) (Figure 4). Community density showed the same trend that species richness: 68 individual plant/m² were found in the 2020s while 40 individual plant/m² were found in the 1980s. However, Shannon diversity was not significantly different between the two periods (2.39 in the 2020s and 2.40 in the 1980s) and the Pielou index was significantly lower in the 2020s (0.69 in the 2020s and 0.78 in the 1980s), demonstrating that the 2020s community were less equitable in abundance (Appendix S1: Figure S4).

**Figure 4.**
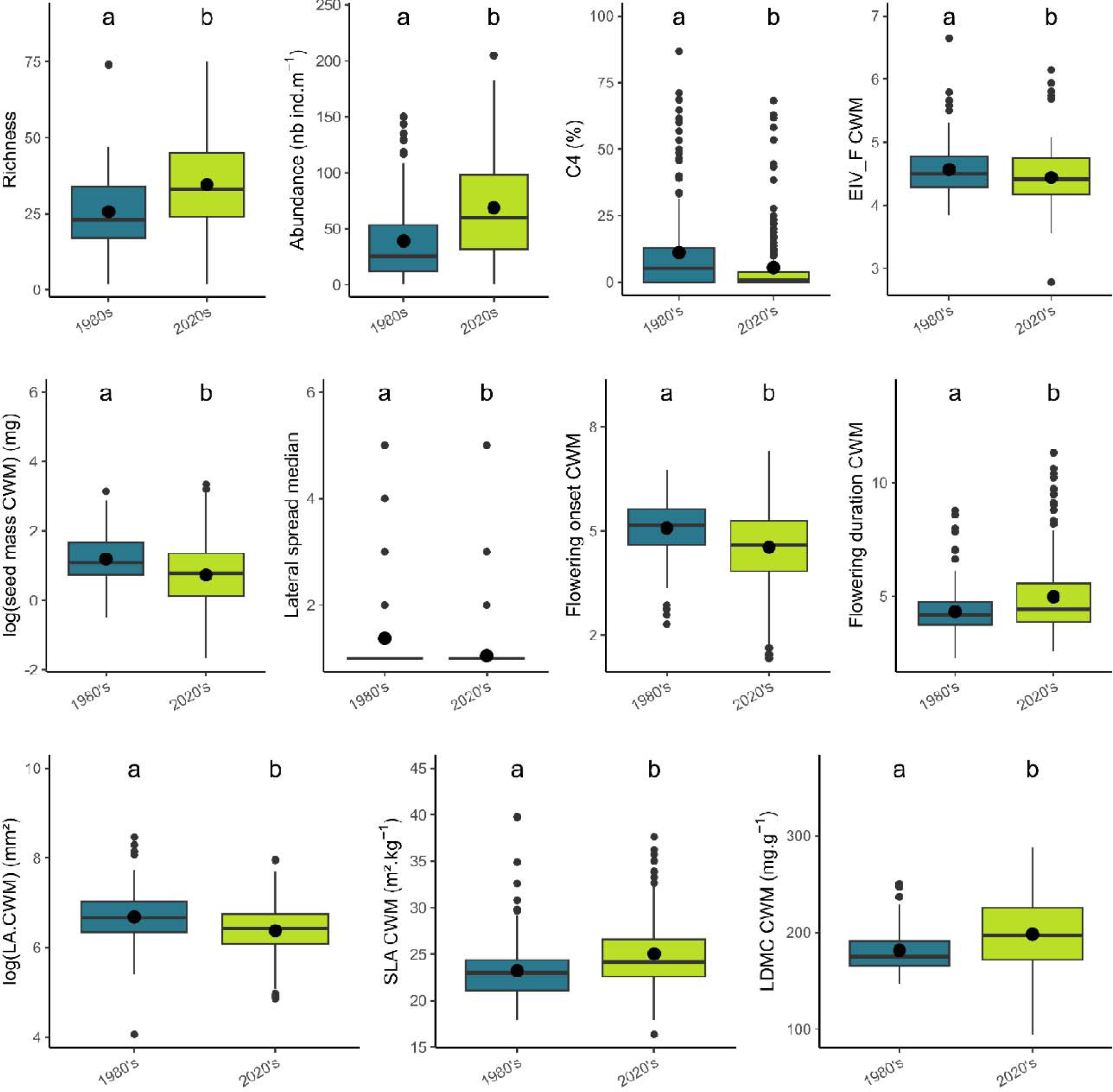
Significant changes in abundance, richness and functional properties of weed communities between the 1980s and the 2020s. CWM, Community Weighted Means; EIV_F.CWM, Ellenberg indice for soil moisture; LA, leaf area; SLA, specific leaf area; LDMC, leaf dry matter content. n = 374 floristic surveys

The 2020s weed communities were composed of more therophytes compared to the 1980s communities (57% of the species in the 1980s compared to 80% in the 2020s) (Appendix S1: Figure S5). In contrast, the 1980s communities were composed of more bulb geophytes (*e.g. Allium vineale*) 7% of bulb geophytes composing 1980s communities while they represented 1% of the weed communities in the 2020s. The 1980s weed communities were composed of more rhizome geophytes (*e.g. Cynodon dactylon*) (10% in the 1980s versus 4% in the 2020s), more biennial hemicryptophytes (*e.g. Chondrilla juncea*) (7% in the 1980s versus 5% in the 2020s), more cespitose and erected hemicryptophytes (*e.g. Anthemis maritima*) (8% in the 1980s and 6% in the 2020s) and more hemicryptophytes with stolons (*e.g. Convolvulus arvensis*) (9% in the 1980s versus 3% in the 2020s).

Out of 423 species, 73 species had a strong decrease in abundance from the 1980s to the 2020s (more than 75%) while 116 species had a strong increase in abundance (more than 75%) (Appendix S1: Figure S6). One hundred and fourteen species strongly decreased in frequency from the 1980s to the 2020s (more than 75%) while 135 species strongly increased (more than +75%) (Appendix S1: Figure S6).

### 3. Shifts in weed community functional structure from the 1980s to the 2020s

Community Weighted Means (CWM) of 13 traits were compared over the two periods (Figure 4). Eight CWM out of 13 were significantly different over the two considered periods. Weed communities from the 2020s had significantly lower leaf area (682 mm² on average in the 2020s and 937 mm² on average in the 1980s), higher SLA (23.3 m².kg^-1^ on average in the 1980s and 25.0 m².kg^-1^ in the 2020s) and LDMC (181 mg.g^-1^ in the 1980s and 198 mg.g^-1^ in the 2020s) (Figure 4). They were also characterised by a higher duration of flowering (4.3 months in the 1980s and 5 months in the 2020s) and earlier flowering species (May on average in the 1980s and mid-April in the 2020s) compared to the 1980s communities. Moreover, the 2020s weed communities were composed of species with lower seed mass (4.2 mg on average in the 1980s and 3.5 mg in the 2020s), lower lateral spreadability, lower soil moisture requirement, and lower proportion of C4 species (11% in the 1980s and 5% in the 2020s).

Over the two periods, weed communities of the Mediterranean vineyard network were mostly ruderal (54% of R-score on average) and competitive (31% of C-score on average) while they had 15% of stress-tolerance-score (Figure 5A). In the 2020s, at the global community pool scale, weed communities were less competitive (27% of C-score) than in the 1980s (34% of C-score) while they were more stress-tolerant (19% of S-score) than in the 1980s (12% of S-score) (Figure 5A, B, C). The R-score was stable over the two periods (54% for both periods) (Figure 5D).

**Figure 5.**
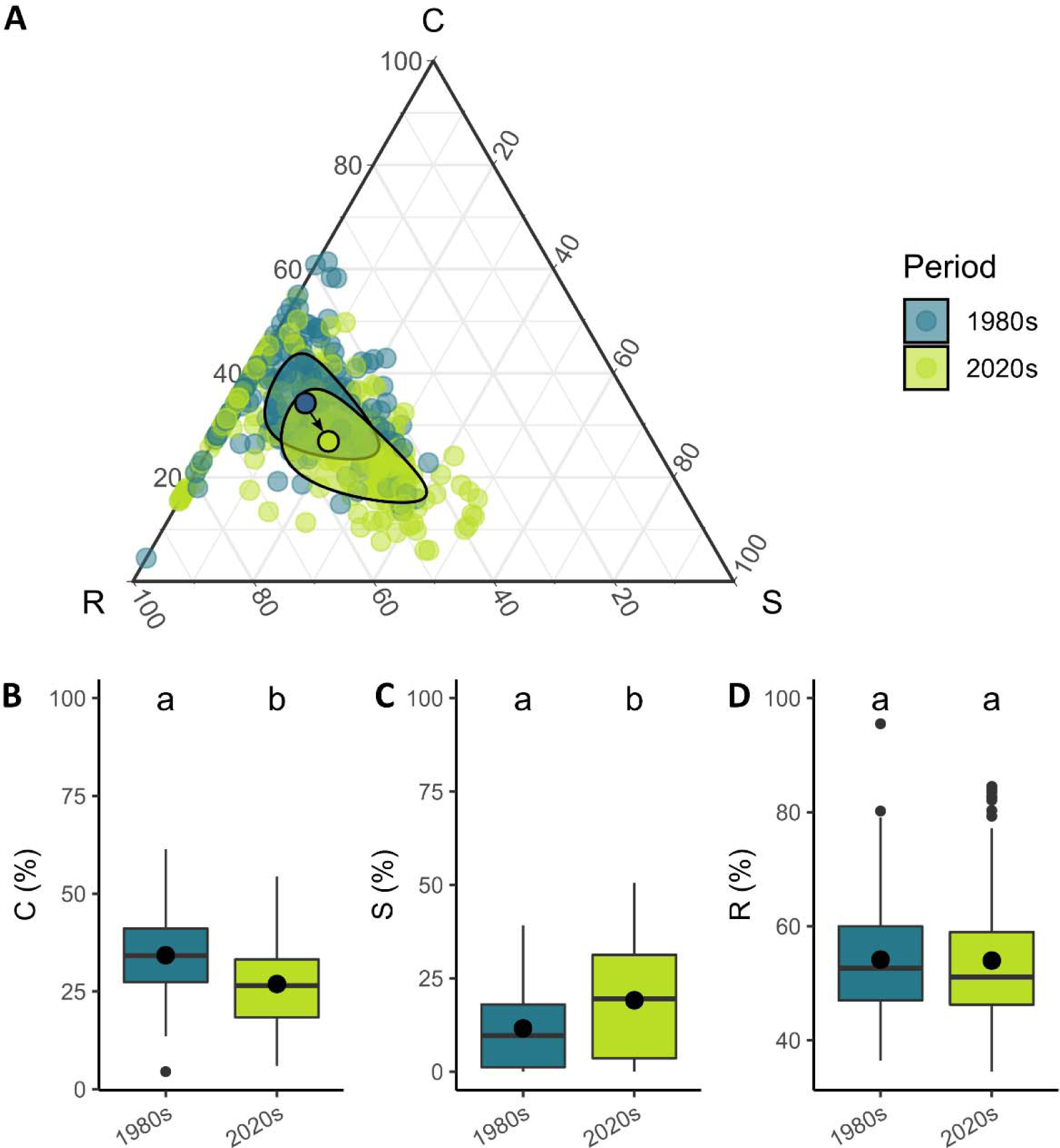
A Shifts in CSR strategies of weed communities from the 1980s to the 2020s in Grime’s triangle. Each dot is a community and their color indicates the period when the community was observed (blue, 1980s; light green, 2020s). The two dots with black outlines are the barycentres of the communities of each period and the black ellipses represent the variance of the CSR strategies of the communities from the two periods. The arrow represents the shift in the mean strategy from the 1980s to the 2020s. Shifts in the proportion of competitive (B), stress-tolerance (C) and ruderal (D) strategies of weed communities from the 1980s to the 2020s. C, Competitors; S, Stress tolerators; R, Ruderal. n = 374 floristic surveys.

Using the elbow inflexion point method on the AUC, the first four dimensions of the CWM space of the 1980s and the 2020s communities were selected (68.9% of total variance explained) (Appendix S1: Figure S7). The first PCA axis opposed late and short-flowering communities to early and long-flowering communities (Appendix S1: Figure S8A, Appendix S2: Table S2). Weed communities of the 2020s were dominated by earlier and longer flowering species compared to the 1980s communities (Appendix S1: Figure S8B, C). The second PCA axis was mainly driven by soil moisture and species temperature requirement (Appendix S1: Figure S8A, Appendix S2: Table S2). Weed communities from the 2020s were composed of species with lower soil moisture requirements and were more thermophilic compared to the 1980s communities (Appendix S1: Figure S8B, D). The third axis of the functional space was mainly driven by the percentage of C4 species within the communities (Appendix S1: Figure S9A). Current weed communities were composed of fewer C4 species compared to the 1980s communities (Appendix S1: Figure S9B, C). Finally, the fourth axis was mostly driven by SLA (Appendix S1: Figure S9A, Appendix S2: Table S2). This axis did not significantly discriminate the 1980s from the 2020s weed communities (Appendix S1: Figure S9B, D). Many CWM correlated significantly (Appendix S2: Table S3). Flowering onset was negatively correlated with the duration of flowering (rho: -0.72, *P*< 0.001) and positively correlated with maximum height (rho: 0.75, *P* < 0.001). The Ellenberg indices for temperature were positively related to Ellenberg indice for light (rho: 0.64, *P* < 0.001) (Appendix S2: Table S3).

### 4. Drivers of the shift in weed community functional and taxonomic structures between the 1980s and the 2020s

Climate change, weed management trajectory from the 1980s to 2020s and the 2020s weed management were not selected in the models explaining changes in abundance and diversity of weed communities between the 1980s and the 2020s. Most of the variance was explained by the vineyard identity and season random effects: 31% for the changes in abundance, 61% for the changes in richness, 27% for the Shannon index and 9% for the Pielou index.

Climate change, the weed management trajectory from the 1980s to the 2020s and the 2020s weed management practices classified into four groups (Chem, Till, Till.IR.Chem.R, Mow.IR.Till.R) were tested as explaining variables of the relative changes in functional properties of weed communities between the 1980s and the 2020s. Vineyards that have experienced a higher climate change intensity from the 1980s to the 2020s (*i.e.* vineyards located in the northern part of the vineyard network, Figure 1) had more shifted to higher LDMC weighted means than vineyards that experienced climate change to a lesser extent (*i.e.* vineyards located nearby the Mediterranean Sea, Figure 1, Table 2). Weed communities that were continuously sprayed with herbicides during the four decades had shifted toward lower CWM of seed mass (-31% on average Appendix S1: Figure S10). Vineyards that were managed toward more integrated weed management (*e.g.* vineyards managed entirely with tillage in the 1980s to vineyard management with mowing, tillage and chemical weeding in the 2020s) were composed of more C4 species in the 2020s than in the 1980s. For the other CWM, the fixed effects were not selected in the final models, after model selection (Appendix S2: Table S4, S5, S6). Weed communities that experienced a higher climate change intensity (*i.e.* higher relative change in annual temperature range over the periods), had shifted toward more stress-tolerance strategies and less ruderal strategies (Table 2). The shifts toward less competitive communities in the 2020s were explained by the adoption of integrative weed management practices: vineyards that were only managed by one type of weed management practice (tillage or chemical weeding) for four decades had less shifted toward a lower score of competitive strategies compared to vineyards where farmers adopted 1 or 2 other types of weed management practices.

**Table 2.**
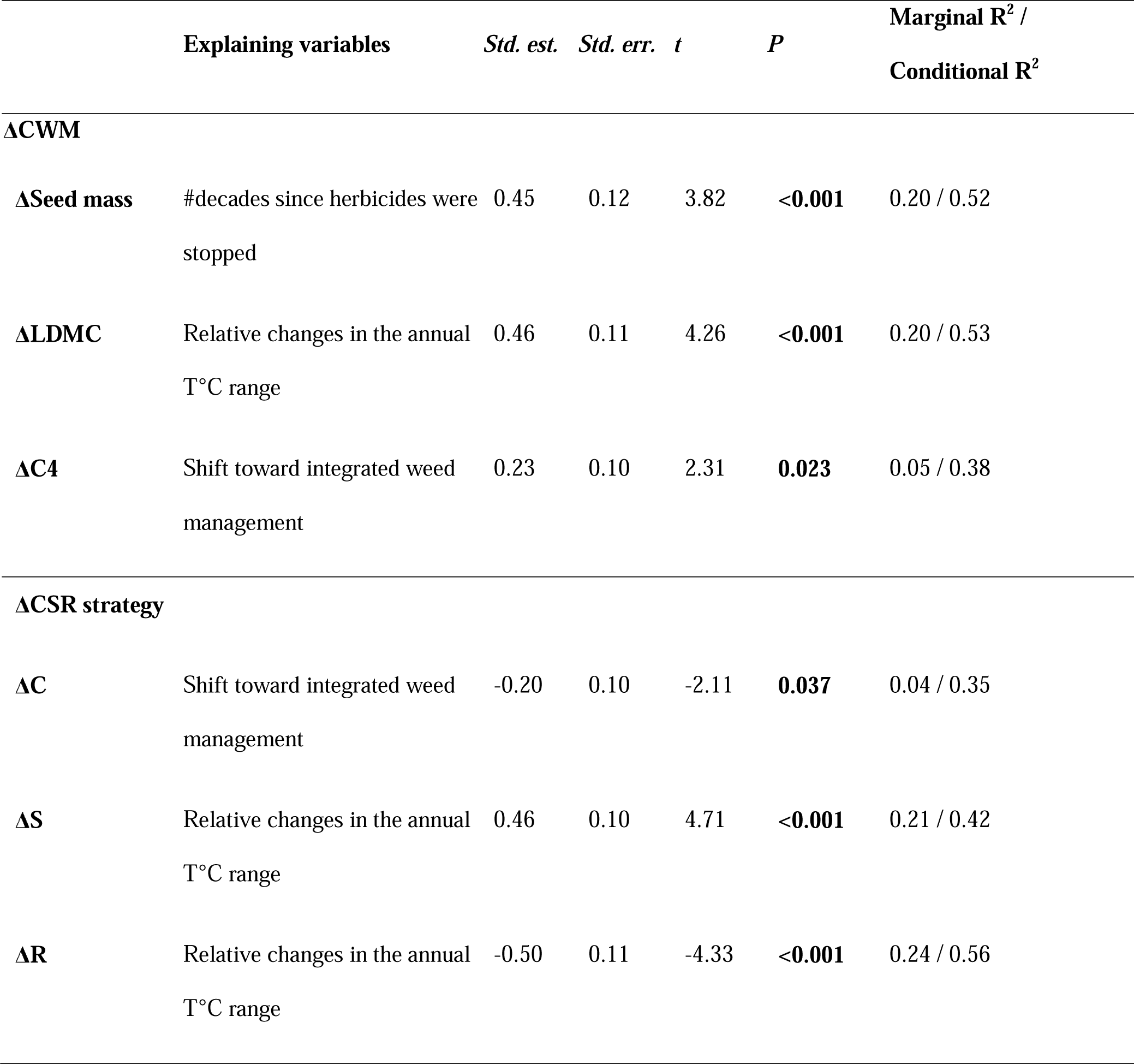
Standardised estimated coefficients of the explaining variables of the relative changes in community weighted means (CWM) and CSR strategies from the 1980s to the 2020s (C, Competitor; S, Stress-tolerators; R, Ruderal). Significant *p*-values (*P* < 0.05) are in bold. n = 108. *Std. est*.; standardised estimates; *Std. err.*, standardised errors; *t*, *t*-value; *P*, *p*-value.

## DISCUSSION

Four main findings were highlighted in this study. First, we demonstrated that climate change occurred during the four decades of the study time scale through the increase of the annual range of temperatures (Figure 6). Second, we found that weed management shifts from the 1980s to the 2020s were mostly characterised by the adoption of mowing and the ending of chemically weeding of the inter-rows. Third, we described major shifts in weed community: (i) higher abundance and richness and lower evenness, (ii) many species increased in abundance and frequency (*e.g. Medicago polymorpha*) while fewer decreased (*e.g. Allium vineale*) and others remained very stable (*e.g. Lolium rigidum*), (iii) community weighted means (CWM) of SLA and LDMC increased, while CWM of leaf area, seed mass, lateral spread and the proportion of C4 decreased, (iv) weed communities shifted toward earlier and longer flowering, lower Ellenberg indice of soil moisture and higher Ellenberg indice of temperature, (v) weed communities shifted toward more stress tolerance strategies and less competitive strategies.

**Figure 6.**
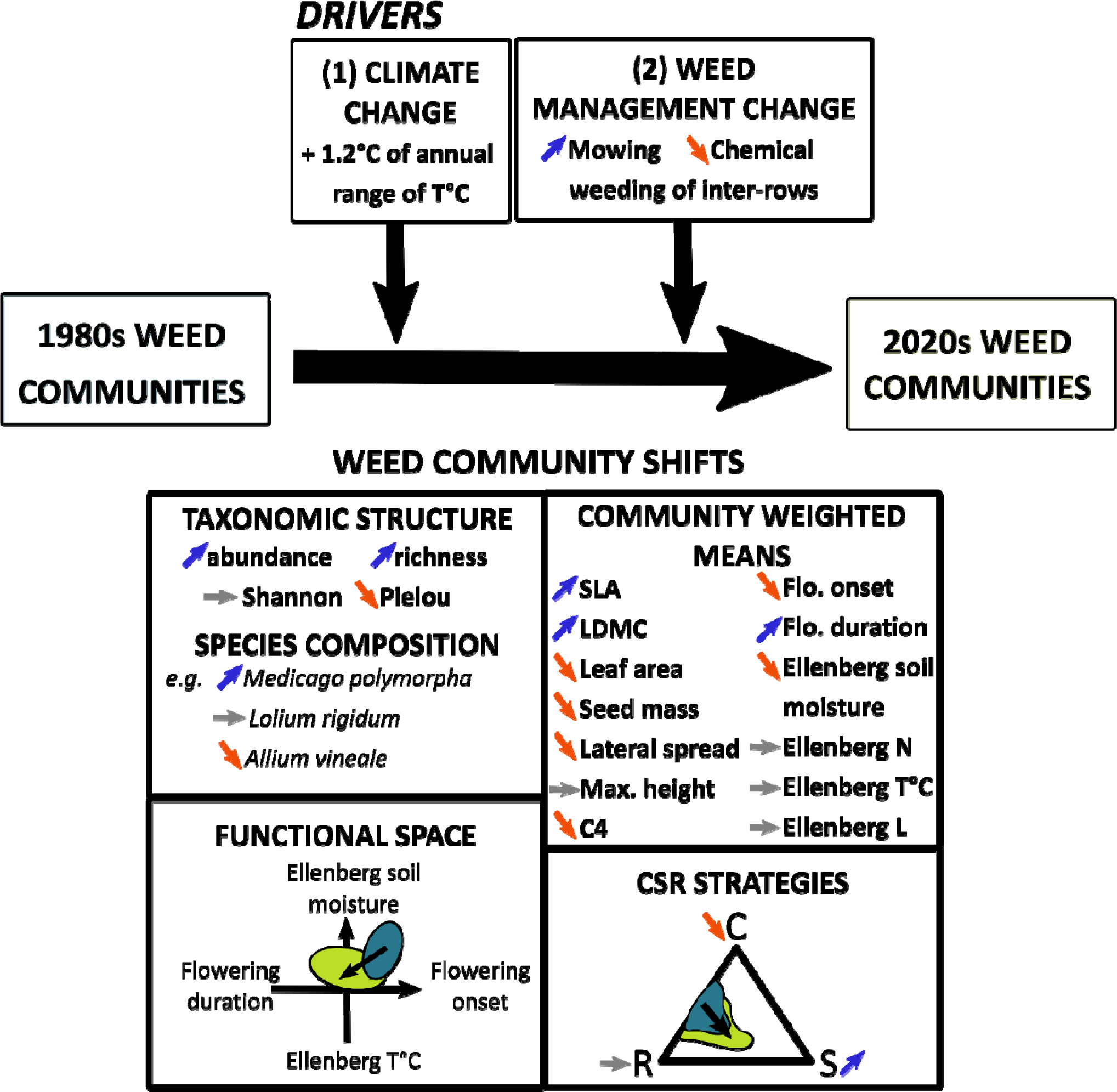
Summary of the main findings of this study. Orange arrows represent significant increases in the variables, blue arrows represent significant decreases and grey arrows represent non-significant changes. Blue areas represent the functional properties of the 1980s communities while green areas represent the functional properties of 2020s communities.

One of the main results of this study is that weed communities from the 2020s were 41% more abundant and 24% more diverse in species than the 1980s communities at the vineyard level. In arable lands, previous studies on weed changes over time showed drastic opposite trends: -42% to -46% in species richness from the 1970s to 2000s (Fried, Dessaint, and Reboud 2016; Fried et al. 2009a) and -67% in species density (Fried et al. 2009b). The contrasts between these studies and our results can be explained by a different temporal coverage. The period 1970s-2000s captured the strong filtering effect of herbicide adoption in arable lands before changes in more extensive weed management practices in the 2000s (Fried et al., 2009a; Richner et al., 2015). On the contrary, the two decades from the 2000s to the 2020s captured the effect of the development of more extensive weed management. However, neither weed management practice trajectory, nor climate change explained the shift in species abundance and richness.

### 1. Significant changes in climate and weed management practices over four decades

Climate shifted from the 1980s to the 2020s toward a higher annual range of temperatures (*i.e.* the difference between the warmest month’s and the coldest month’s mean temperatures) that averaged around 1.2 °C. Seasonal temperature fluctuations of climate are expected to occur under climate change (Peters, Breitsameter, and Gerowitt 2014) and annual variability of temperatures was the best predictor of climate change in our dataset. While the studies of climate change in the Mediterranean area projected a decrease in precipitations (Ali et al. 2022), seasonal precipitation variability over time explained fewer climate shifts than temperature metrics from the 1980s to the 2020s. Precipitation variability within the vineyard network was greater than its variability over time.

We hypothesised that weed management practices would evolve toward less chemical weeding use and more frequent tillage and mowing from the 1980s to the 2020s (Cataldo, Fucile, and Mattii 2021). Globally, weed management shifts matched our expectations and were driven by the adoption of mowing of the inter-rows that replaced chemical weeding of the inter-rows. More surprisingly, chemical weeding of rows and inter-rows was not widespread in 1978 and 1979 (*i.e.* the years of the first floristic surveys of this study) in the vineyard network. Indeed, 14 vineyards were managed using herbicides on both rows and inter-rows on 40 vineyards. Tillage of rows and inter-rows was still quite commonly applied (9/40 vineyards). Chemical weeding spread more widely in the decade that followed the end of the 1970s, a few years after the first floristic surveys of this study. Maillet (1992) quantified for an extended network of vineyards around Montpellier (that included the vineyard network of this study) that 41% of vineyards shifted towards chemical weeding of both rows and inter-rows from 1979 to 1987. Moreover, in the 1990s, the frequency of chemical weeding globally increased with a second application at the end of spring to control for summer species (Maillet 1992). In our study, we did not consider the shift in frequency of weed management and herbicide dose due to the difficulty of obtaining this information from farmers for the decades between the 1980s and 2020s. Thus, if the median management that was used in the 1980s and the 1990s is the same in our management trajectory characterisation, it is most likely that 1990s weed management was more intensive in herbicide use. We also did not explicitly consider in the characterisation of the weed management changes, the active ingredients of herbicides that were different between the two periods. In the 1980s, 95% of herbicides used were simazine (pre-emergence) and aminotriazole-based (post-emergence) (Maillet 1981). In the 2020s, herbicides were glyphosate-based (post-emergence herbicide). If the spectrum of these herbicides from both periods was broad (*i.e.* targeting both annuals and perennials), persistence over time was different: simazine can persist from three months to 15 months while aminotriazole and glyphosate persist less than one month (Maillet 1981). Thus, in the 1980s, chemical weeding had a longer controlling action on weed communities. This could explain for instance, that even the plots that were continuously chemically weeded between the 1980s and the 2020s showed a slight increase of species richness (13%).

### 2. Climate change-induced weed community shifts toward more stress-tolerance

According to our hypothesis, at the regional scale, weed communities shifted toward more stress tolerance (+37% for S-score). Weed communities, that shifted toward a higher score of stress tolerance from the 1980s to the 2020s, have also experienced a higher intensity of climate change (*i.e.* a higher increase in annual temperature range). To our knowledge, this study is the first to demonstrate that stress tolerance of weed communities is increasing in response to climate change through changes in species composition.

The increase in stress tolerance is linked to the increase of the community-level leaf dry matter content (181 mg.g^-1^ in the 1980s and 198 mg.g^-1^ in the 2020s) which is the *proxy* for stress tolerance in Pierce et al.’s algorithm (2017). The increase in LDMC was also explained by the increase in annual range temperatures in the selected models. This indicates that climate change intensity reshuffled community composition toward more conservative species which invest in leaves with thicker and stiffer cell walls that can maintain turgidity at low water potential (Garnier et al. 2019). Other shifts of community-level traits indicated more conservative and stress tolerance communities even though they were not significantly linked to the intensity of climate change. For instance, leaf area decreased over time which has been related to a drought-tolerance strategy (de la Riva et al. 2018). Ellenberg’s indice of soil moisture of weed communities also decreased significantly. Weed communities had earlier and longer flowering times in the 2020s compared to the 1980s which could also be explained by a longer growing season (Menzel et al. 2006). Surprisingly, the proportion of C4 species within the weed communities decreased over time while it is expected that C4 species would increase with climate change as their photosynthesis pathway is more efficient than those of C3 at higher temperatures (Peters, Breitsameter, and Gerowitt 2014; Heilmeier 2019). However, independent of temperature and precipitation changes, the increasing level of CO2, which we did not consider, has been predicted to favour C3 more than C4 (Peters, Breitsameter, and Gerowitt 2014). Indeed, C3 weeds increase their leaf area and their biomass in higher CO2 pressure than C4 species. Thus, they are more competitive in such conditions.

### 3. The diversification of weed management practices over time favoured less competitive weed communities

We hypothesised that the decrease in herbicide use replaced by tillage and mowing would favour more conservative and competitive strategies for two reasons. First, mowing is a lower-intensity disturbance than chemical weeding and would filter more conservative strategies than other management practices (MacLaren, Bennett, and Dehnen-Schmutz 2019). Second, weed biomass and competition for resources might be higher in mown communities, selecting competitive species. In contrast with our expectations, communities from the 2020s were less competitive than communities from the 1980s (34% of C-score in the 1980s versus 27% of C-scores in the 2020s) while mowing appeared in the 2000s in addition to the frequent tillage practice, resulting in more complex weed management. A shift toward a higher diversity of weed management practices and consequently more integrated management of weeds induced a higher decrease in C-score. This strategy score is determined by leaf area in Pierce et al.’s algorithm (2017) which also decreased over time. The diversity of disturbance types, intensity and frequency of integrated managed vineyards favoured communities composed of species that did not invest in large leaf areas and high maximum height (which was positively correlated to leaf area in our study) and thus were less competitive to capture light. However, this was in contradiction with the higher SLA of weed communities from the 2020s compared to the 1980s. Even though the effect of integrated weed management on C-score was quite low (estimate: -0.20) and with a high *P*-value (0.037), these results demonstrate that weed management is a lever of action to filter more desirable weed communities that are less competitive with the main crops (MacLaren et al. 2020).

Weed communities continuously sprayed with chemical weeding for four decades shifted toward lower seed mass (on average, -31% of seed mass at the community level) while those where herbicides were interrupted for more than 20 years, showed the opposite trend (+22% of seed mass). As seed mass is negatively correlated to seed number (Westoby 1998), weed community strategies under chemical control are to produce a high number of seeds to face a higher risk of seed mortality (Storkey, Moss, and Cussans 2010). Moreover, small seeds may have higher seed dormancy (Thompson, Band, and Hodgson 1993) leading to a lower depletion of the seed bank. In tilled vineyards, seeds can be buried in the top 5 centimetres of the soil after tillage and are less exposed to desiccation. As heavier seeds are most likely to germinate when they are buried (Benvenuti, Macchia, and Miele 2001), the weed community strategy under tillage is to produce large seeds with a better probability of seedling establishment.

### 4. Future weed communities: a threat or an opportunity for vineyards in the context of climate change?

The trend toward more stress-tolerant, less competitive and more diverse weed communities that we identified for perennial cropping systems contrasted with several studies based on annual cropping systems (Storkey, Moss, and Cussans 2010; Moss et al. 2004). In annual cropping systems, the temporal increase in weed competitiveness was observed from 1969 to 2014 in the Broadbalk Experiment, UK and was partly related to an increase in nitrogen input (Storkey et al. 2021). Fertilisation is a factor that we did not consider in our study management due to the difficulty of obtaining this data from farmers. However, fertilisation amount in Mediterranean vineyards is quite low (Metay et al. 2014) and the global temporal trend of fertilisation in vineyards contrasted with the trend in annual cropping systems. Indeed, vineyard fertilisation has decreased over the past decades with the aim of increasing wine quality (Verdenal et al. 2021) and this trend could also explain the decrease in weed community competitiveness. Weed management is a way for farmers to manage temporal variations in vine water supply which is a key determining factor of crop productivity and fruit quality (Leeuwen et al. 2009). Changes in weed management strategies (*e.g.* choice to sow a cover crop in the inter-rows) and weed management tactics (*e.g.* choice of the date of weed destruction) are both needed to adapt to higher temporal uncertainty, for short and long-term changes.

## Conclusion

Using weed communities in Mediterranean vineyards as a model, we showed that management and climate changes are modelling plant communities over time. The current shift towards more extensive weed management in vineyards is driving more diverse and abundant plant communities. In response to climate change, weed communities were more stress-tolerant. The diversification of weed management techniques based on less herbicide use and more tillage or mowing practices, at the plot level, drove less competitive weed communities. We showed that the CSR framework is a relevant framework to assess changes in environmental conditions along the resource gradient and shifts in management along the disturbance gradient. In this article, we demonstrated that plant communities are adapting to climate change and that land management is a strong lever for action to model diverse and functional plant communities.

## Acknowledgements

This work was supported by Occitanie Region [Arrêté modificatif N° 19008795 / ALDOCT-000660 Subvention d’investissement, Allocations de recherche doctorales 2019] and the Office Français de la Biodiversité [ECOPHYTO II: Axe 2 – Action 8 and 9, N°SIREPA: 4148] as part of the SAVING project: Spatio-temporal dynamics of weed species communities in response to soil management practices in vineyards and consequences for grapevines: transition to zero glyphosate management. Our work was also partly funded by the French National Research Agency (grant ANR-16-CE02-0007). We thank all winegrowers who provided management information and access to their farms. Many thanks to Manon Alvanitakis, Amélie Horain, Anna Orvoire, Victor Berteloot, Isis Poinas and Simon Poulet for their help in the field. We thank Eric Garnier for his assistance in digitizing data, Taïna Lemoine for her kind help in R and Maxime Henriquet for his assistance in QGis.

## Author contributions

Marie-Claude Quidoz formatted the database. Marie-Charlotte Bopp led the writing of the manuscript. All authors contributed critically to the drafts and gave final approval for publication.

