## Appendix S1 for "Climate and management changes drove more stress-tolerant and less competitive plant communities in 40 years"

### Appendix 1

### Title

Climate and management changes drove more stress-tolerant and less competitive plant communities in 40 years

**Authors**

Marie-Charlotte Bopp^1*^, Elena Kazakou^1^, Aurélie Metay^2^, Jacques Maillet^3^, Marie-Claude Quidoz^1^, Léa Genty^2,5^, Guillaume Fried^4^

**Ecological monographs**

### Supporting figures

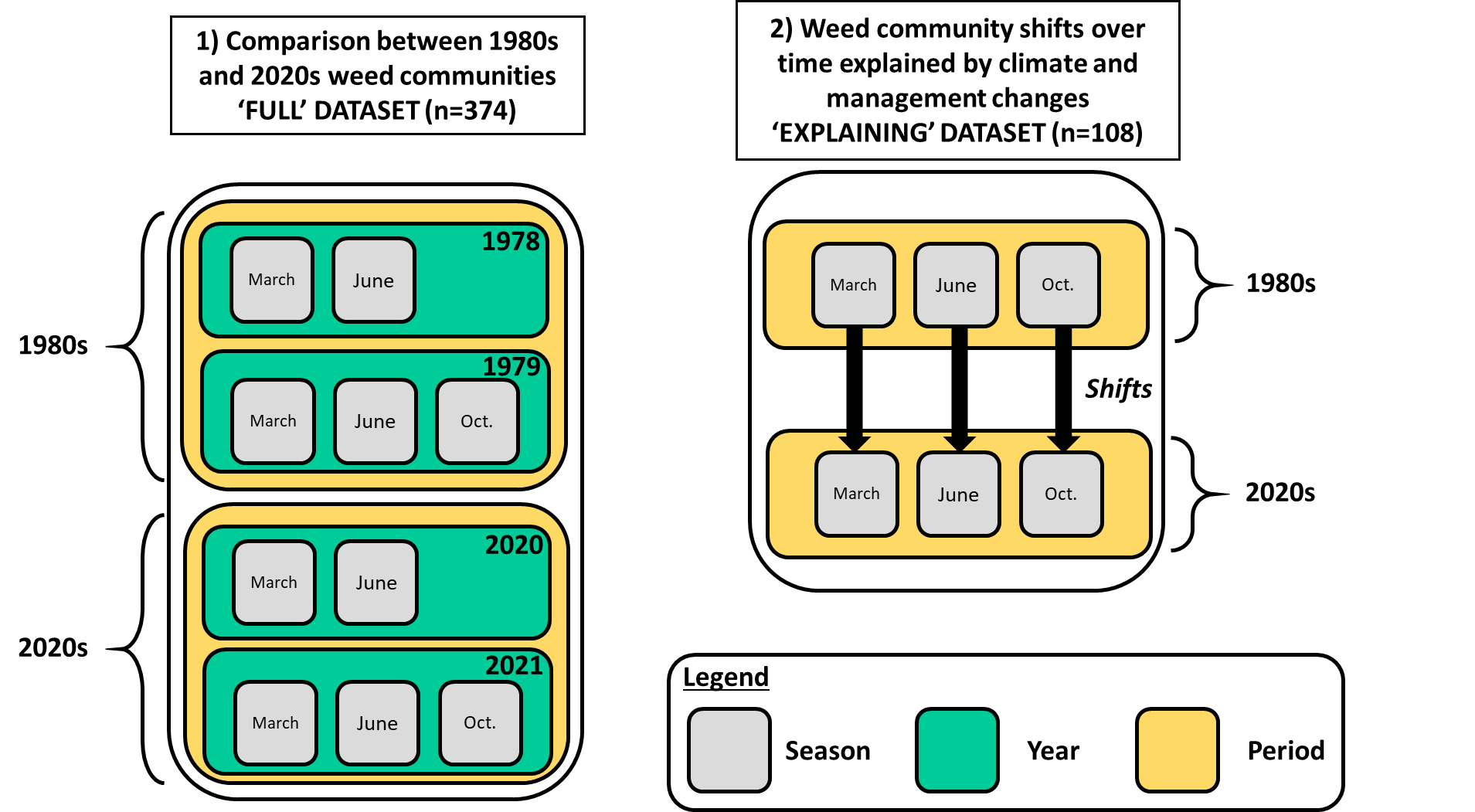

Figure S1 Two different datasets were used for data analyses. The first one (on the left) is the ‘full dataset’ composed of 374 floristic surveys from the 1980s and the 2020s, with all seasons (March, June, Oct. ; October) and years (1978, 1979, 2020, 2021). The second dataset (on the right) is the ‘explaining dataset’ composed of 108 weed community shifts of the same seasons from the 1980s to the 2020s. Because of a summer drought in 1978, the autumn floristic surveys were not done that year (Maillet 1981). Therefore, to have equivalent data sets for the 1980s and 2020s periods, we did not use the floristic surveys of Autumn 2020. Moreover, in March 1978 and March 2020, only 27 plots were surveyed. In total, the data set was composed of 374 floristic surveys (2 periods x 27 vineyards x 1 season + 2 periods x 40 vineyards x 4 complete seasons).

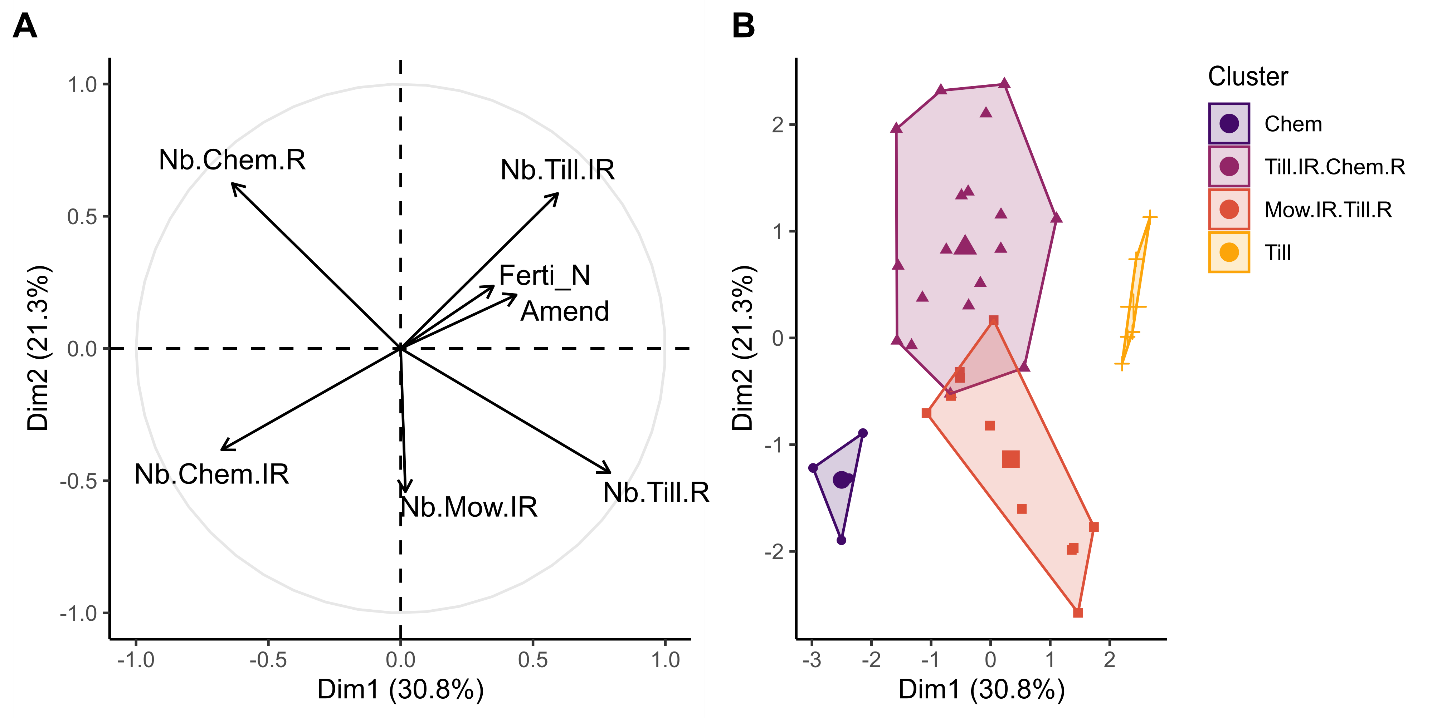

Figure S2**A** Circle of correlation of the agricultural practices applied by the farmers from 2015 to 2021 projected on the first two axes of the Principal Component Analysis (PCA) (Dim1, Dim2). Each practice variable is averaged over the 2015-2021 period. Nb.Chem.IR, number of chemical weeding sprayed in the inter-rows ; Nb.Chem.R, number of chemical weeding sprayed in the rows ; Nb.Till.IR, number of tillage of the inter-rows ; Nb.Till.R, number of tillage of the rows ; Nb.Mow.IR, number of mowing of the inter-rows ; Ferti_N, nitrogen inputs ; Amend, soil amendment. **B** Clusters of vineyards sharing similar agricultural practices displayed on the first two axes of the PCA. Each dot is a vineyard and its colour indicates to which cluster it belongs. Vineyards were classified in four clusters. Cluster ‘Chem’ gathers four vineyards with chemical weeding of both rows and inter-rows. Cluster ‘Till.IR.Chem.R’ regroups 19 vineyards with tilled inter-rows and chemically weeded rows. Cluster ‘Mow.IR.Till.R’ gathers 11 vineyards with mowed inter-rows and tilled rows. Cluster ‘Till’ groups six vineyards with tilled rows and inter-rows.

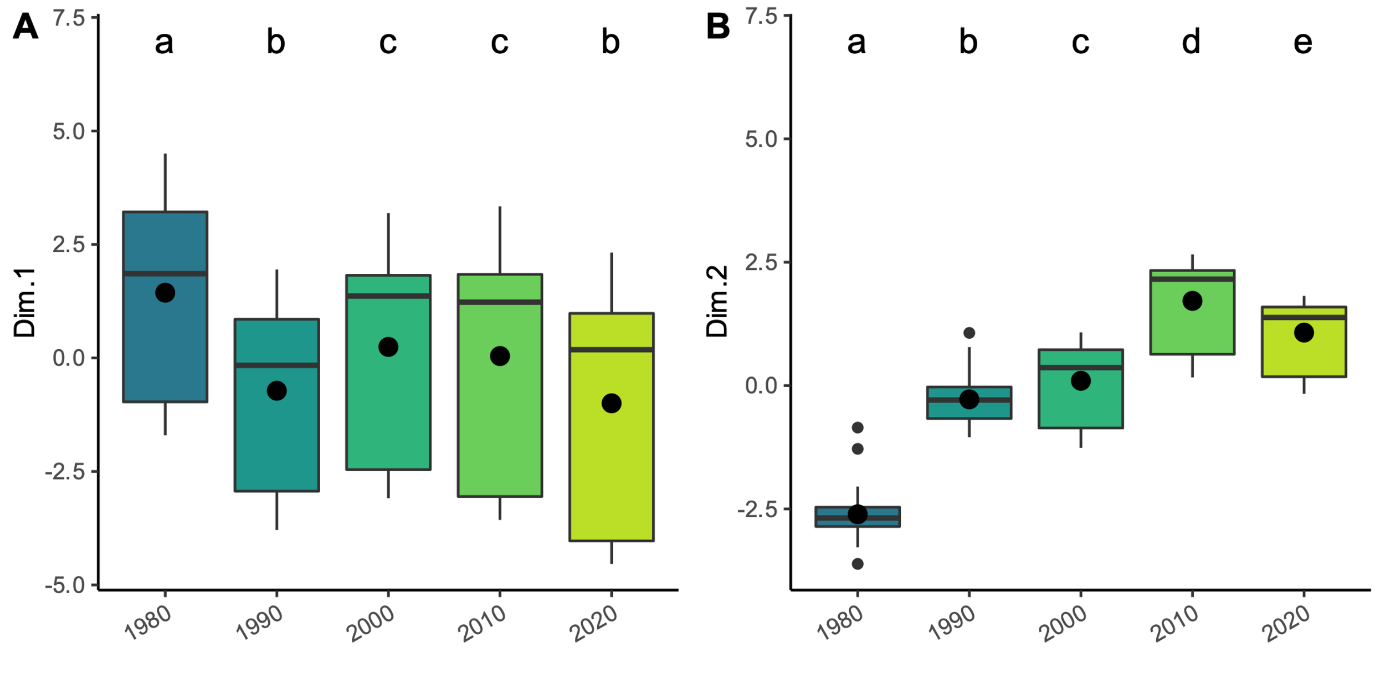

Figure S3 Climate PCA coordinate changes of vineyards on the first axis (A) and the second axis (B) over the decades from the 1980s to the 2020s.

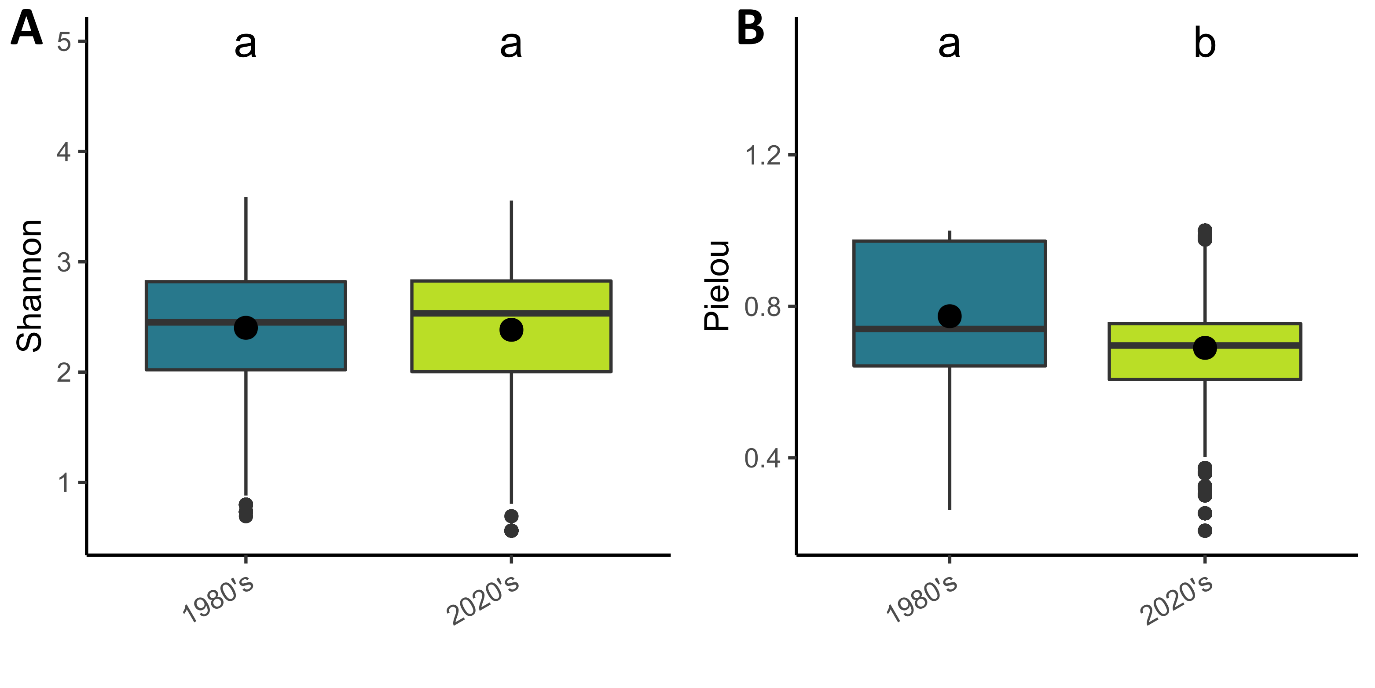
Figure S4 Significant changes in Shannon and Pielou indices of weed communities between the 1980s and the 2020s. n = 374 floristic survey.

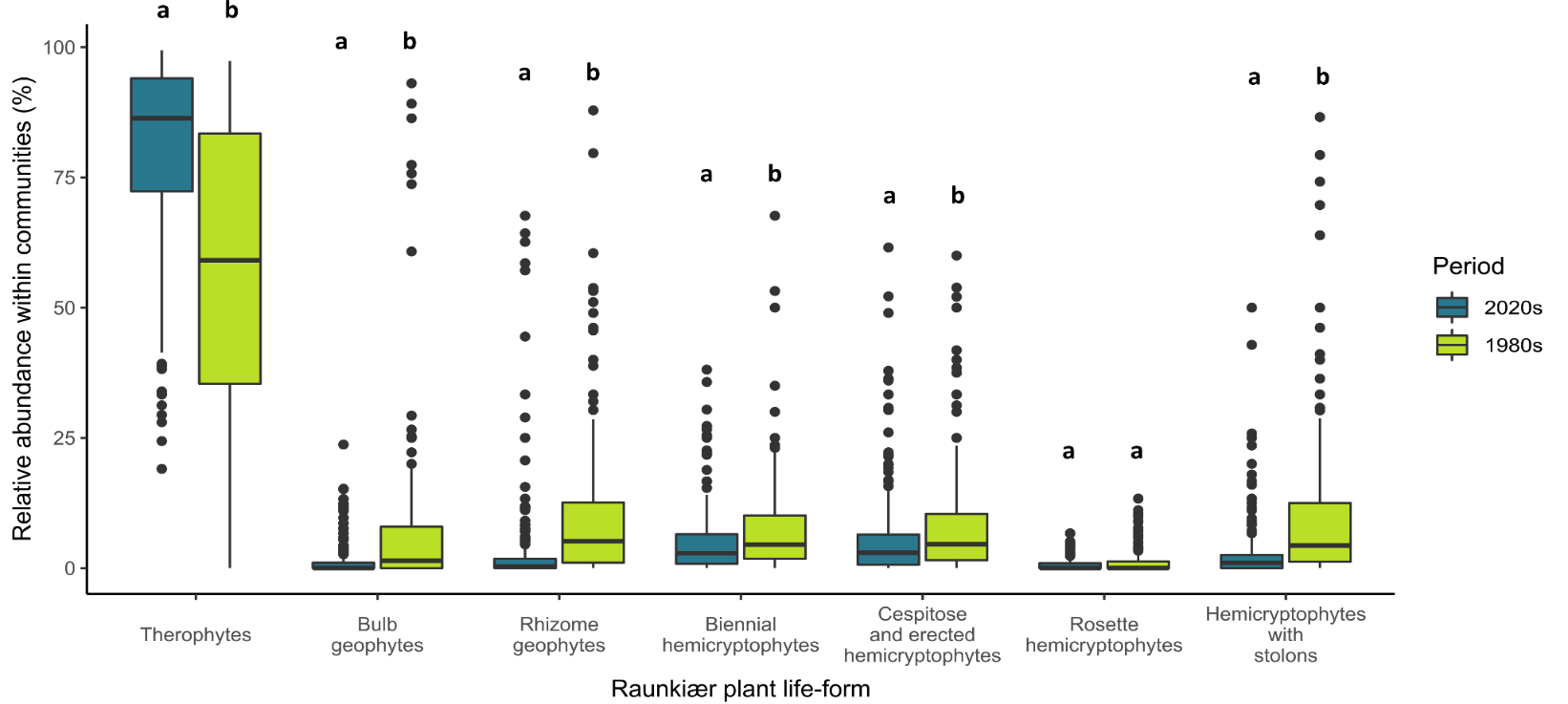

Figure S5 Relative abundance of Raunkiær plant-life forms within the 1980s and 2020s weed communities. n = 374 floristic survey.

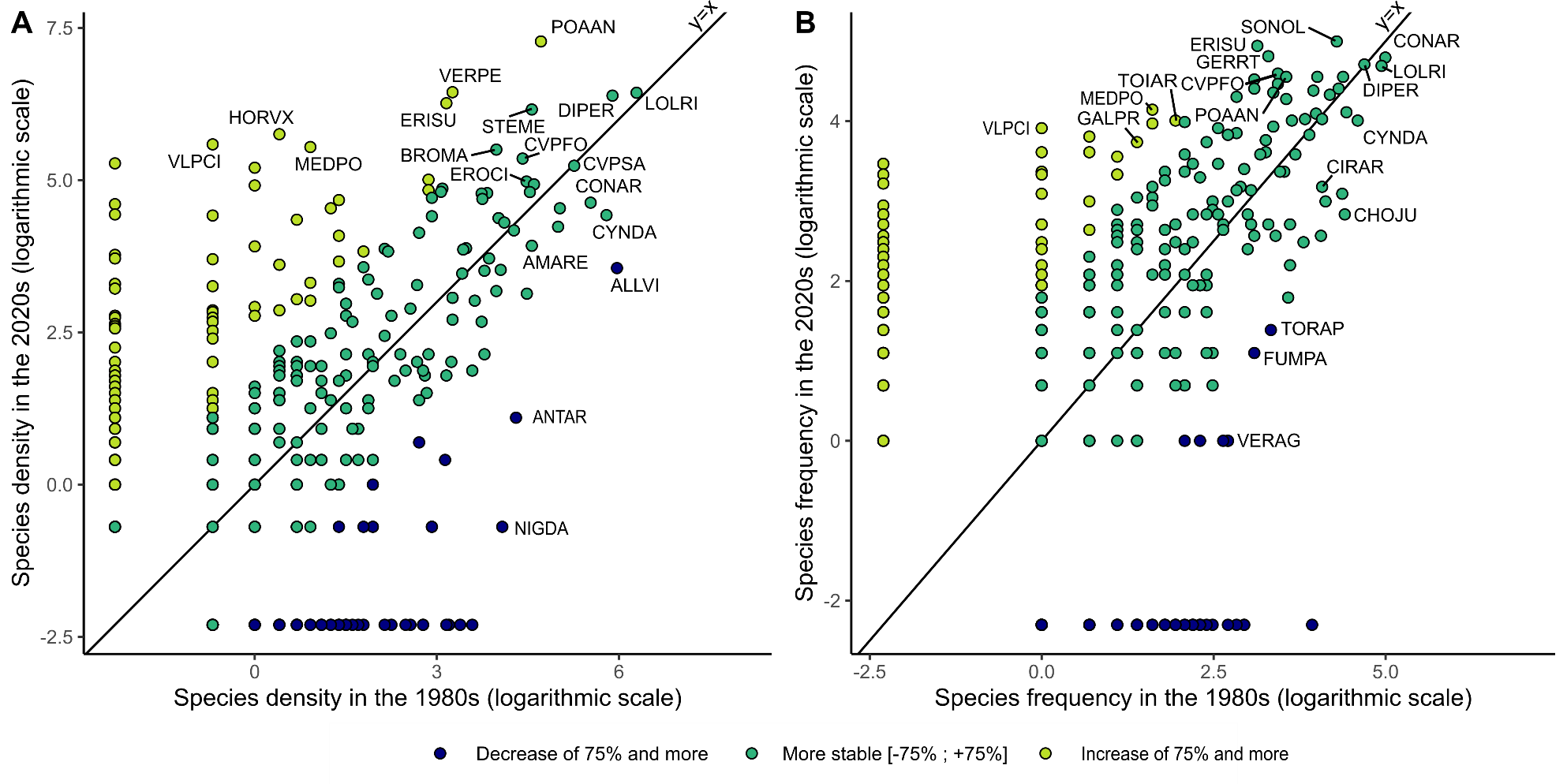

Figure S6 Changes in density (A) and frequency (B) for every 423 species that were found in the 1980s and/or in the 2020s. Each dot is a species and its colour describes to which extent its density and frequency have changed after four decades: dark blue dots represent species with density or frequency that decreased more than 75% from the 1980s to the 2020s; light green dots are the species with density or frequency that have increased by 75% or more; green dots are the more stable species. The dark line represents the y=x line. The most abundant and frequent species that decreased the most, that increased the most or that remained stable are identified. Among the most abundant species in the 1980s and/or 2020s, the density of *Poa annua* (+86% of increase), *Veronica persica* (+92%), *Erigeron sumatrensis* (+91%), *Vulpia ciliata* (+99%) and *Medicago polymorpha* (+98%) had strongly increased during the last 40 years. In contrast, the density of *Allium vineale* (-84%), *Cynodon dactylon* (-59%), *Cirsium arvense* (-59%), *Anthemis arvensis* (-92%) and *Nigella damascena* (-98%) had strongly decreased over time. *Lolium rigidum* (+7%) and *Crepis sancta* (-1%) were among the most stable and abundant species in abundance over time. Among the most frequent species in the 1980s and/or 2020s, the frequency of *Medicago polymorpha* (+85% of increase), *Torilis arvensis* (+77%), *Galium parisience* (+83%) and *Vulpia ciliata* (+96%) increased strongly over time. In contrast, the frequency of *Tordylium apulum* (-75%), *Fumaria parviflora* (-76%) and *Veronica agrestis* (-88%) strongly decreased. *Lolium rigidum* (-12%) and *Diplotaxis erucoides* (+1%) were among the most stable and the most frequent species over time in the network. ALLVI, *Allium vineale* ; AMARE, *Amaranthus retroflexus* ; ANTAR, *Anthemis arvensis* ; BROMA, *Anisantha madritensis* ; CHOJU, *Chondrilla juncea* ; CIRAR, *Cirsium arvense* ; CONAR, *Convolvulus arvensis* ; CVPFO, *Crepis foetida* ; CVPSA ; *Crepis sancta* ; CYNDA, *Cynodon dactylon* ; DIPER, *Diplotaxis erucoides* ; ERISU, Erigeron sumatrensis ; EROCI, Erodium cicutarium ; FUMPA, Fumaria parviflora ; GALPR, *Galium parisience* ; GERRT, *Geranium rotundifolium* ; HORVX, *Hordeum vulgare* ; LOLRI, *Lolium rigidum* ; MEDPO, *Medicago polymorpha* ; NIGDA, *Nigella damascena* ; POAAN, *Poa annua* ; SONOL, *Sonchus oleraceus* ; STEME, *Stellaria media* ; TOIAR, *Torilis arvensis* ; TORAP, *Tordylium apulum* ; VERAG, *Veronica agrestis* ; VERPE, *Veronica persica* ; VLPCI, *Vulpia ciliata*.

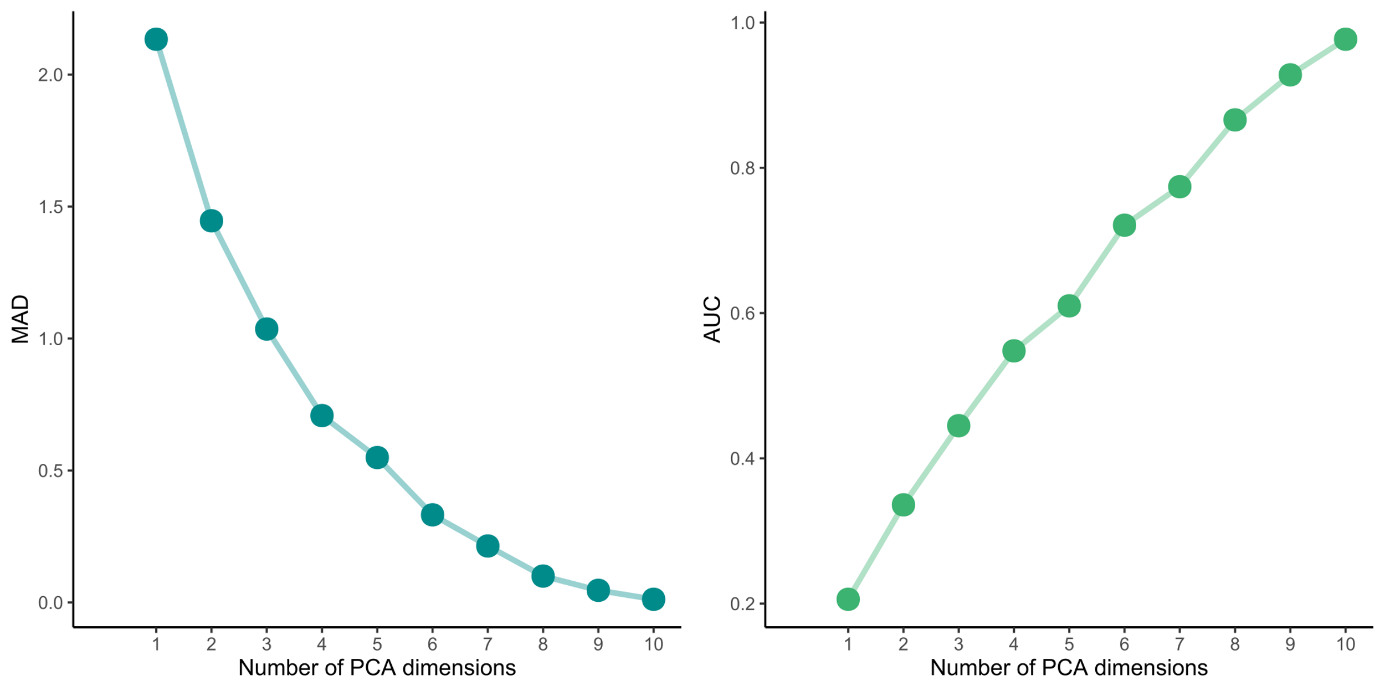

Figure S7 The quality of the community functional space according to the selected number of PCA dimensions: (i) the AUC (Area Under the Curve) based on the Somer’s D statistic which measures the difference of ranking between the communities of the initial data set and the communities displayed in the PCA and (ii) the MAD (Mean Absolute Deviation) measures the Euclidean distance between communities’ position in the initial data set and the PCA. Using the method of the elbow infection point method proposed by Mouillot et al. (2021) on AUC, we determined that 4 dimensions would be necessary to describe the functional space well.

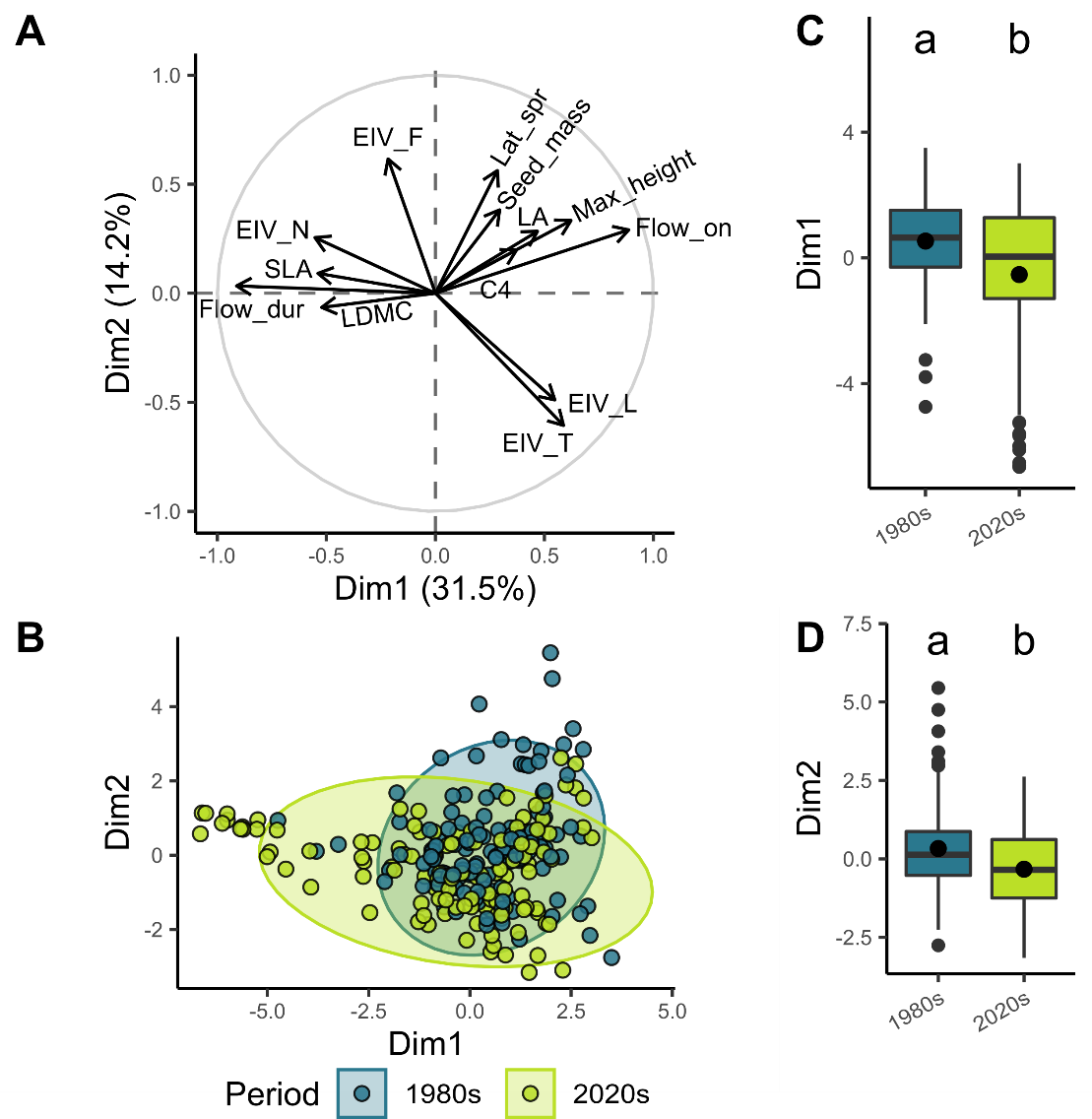

Figure S8 Functional shift in the weed communities from the 1980s to the 2020s along the first two axes of the PCA based on community-weighted means (CWM). A) CWM constructing the first two axes of the functional PCA are displayed in the correlation circles. B) The weed communities are projected on the first two dimensions. Each dot is a weed community and its colour refers to the period in which weed communities were observed: blue, 1980s ; green, 2020s. The significance of the differences in the coordinates of weed communities from the 1980s and the 2020s is presented for the first (C) and the second (D) dimensions of the functional PCA. n = 374 floristic survey.

Figure S9 Functional shift in the weed communities from the 1980s to the 2020s along the third and fourth axes of the PCA based on community weighted means (CWM). A) CWM constructing the third and fourth axes of the functional PCA are displayed in the correlation circles. Poorly represented CWM were removed from the graph for reasons of readability (flowering onset and flowering duration). B) The weed communities are projected on the third and fourth axes. Each dot is a weed community and its colour refers to the period in which weed communities were observed: blue, 1980s ; green, 2020s. The significance of the differences in the coordinates of weed communities from the 1980s and the 2020s are presented for the third (C) and the fourth (D) dimensions of the functional PCA. n = 374 floristic survey.

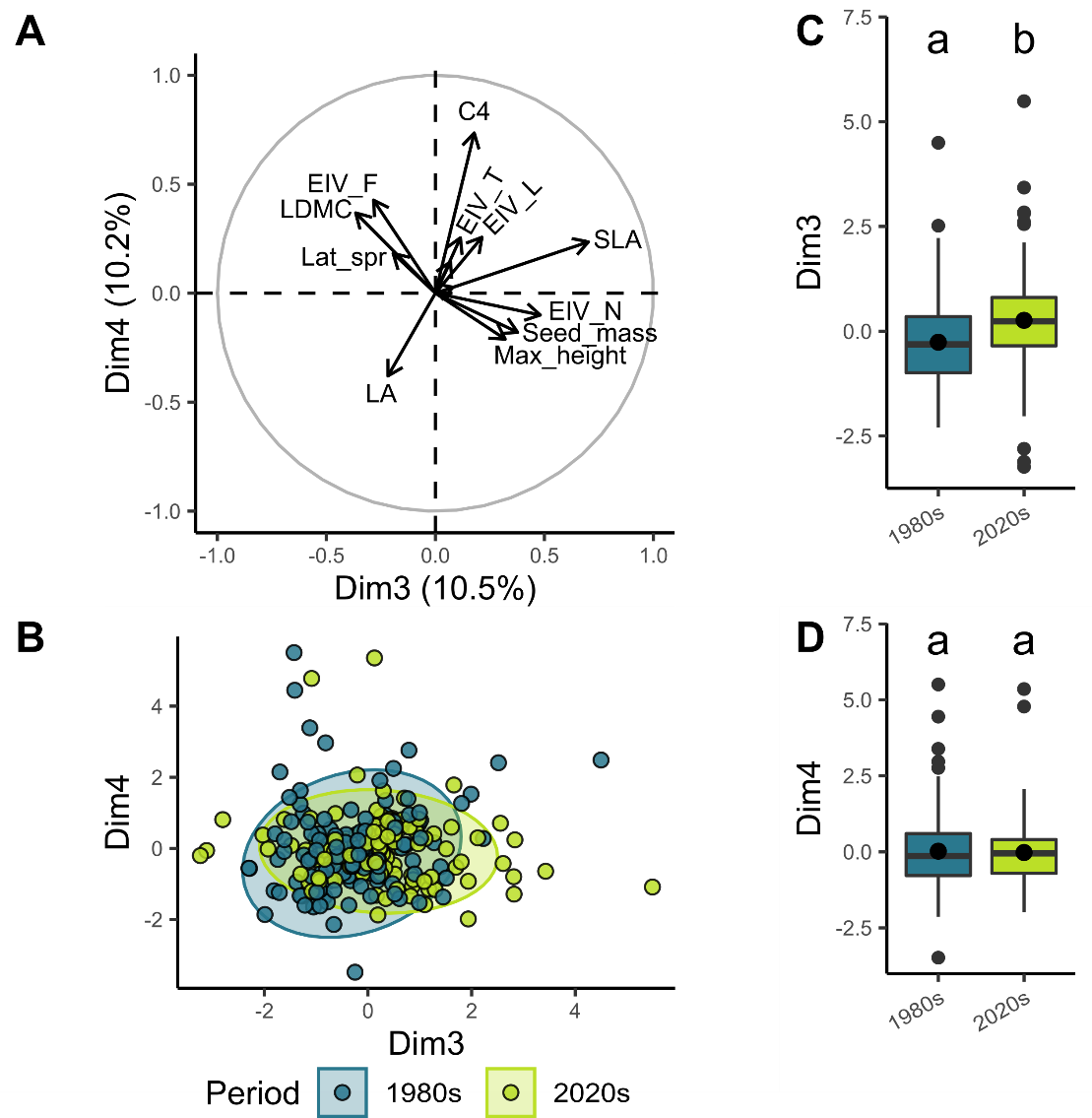

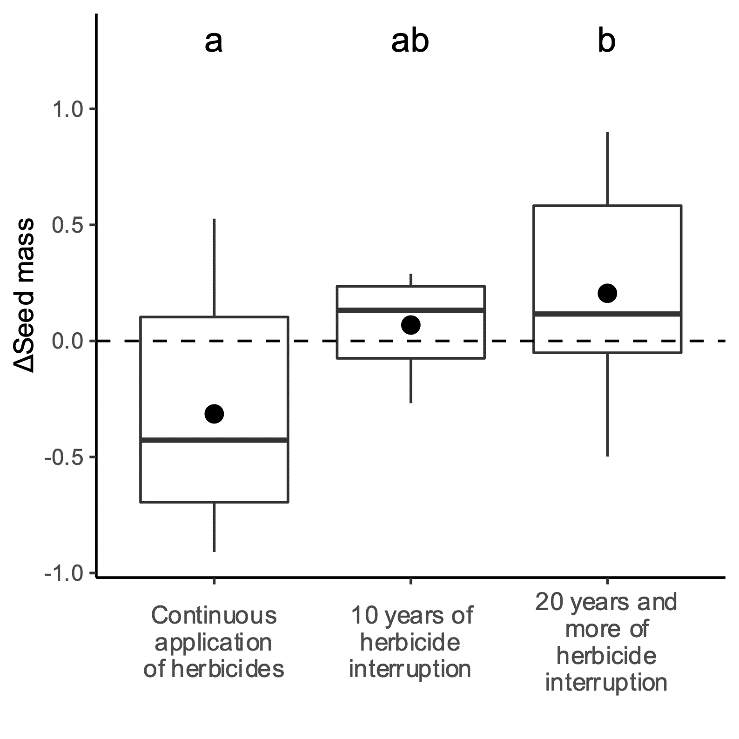

Figure S10 Seed mass shift according to number of decades since herbicides were stopped. ΔSeed mass is equal to (Community weighted seed mass2020s – Community weighted seed mass1980s) / (Community weighted seed mass2020s + Community weighted seed mass1980s). n = 108.
