## Appendix S2 for "Climate and management changes drove more stress-tolerant and less competitive plant communities in 40 years"

### Appendix 2

### Title

Climate and management changes drove more stress-tolerant and less competitive plant communities in 40 years

**Authors**

Marie-Charlotte Bopp^1*^, Elena Kazakou^1^, Aurélie Metay^2^, Jacques Maillet^3^, Marie-Claude Quidoz^1^, Léa Genty^2,5^, Guillaume Fried^4^

**Ecological monographs**

### Supporting tables

| Table S1 Description of the metrics used to evaluate the shifts in climate, weed management, functional and taxonomic structures of the weed communities. | | | |
| --- | --- | --- | --- |
| **Type of drivers** | | **Variable description** | **Computed metric** |
| Climate change intensity | | Relative change in the annual temperature range | $\Delta T^{\circ}C=\frac{{T^{\circ}C range}_{2020's}-{T^{\circ}C range}_{1980's}}{{T^{\circ}C range}_{2020's}+{T^{\circ}C range}_{1980's}}$  With$T^{\circ}C range$ which is the difference between the warmest month's and the coldest month's mean temperature |
| Weed management trajectory from the 1980s to the 2020s | | The number of decades since the application of herbicides was stopped |  |
|  |  | The number of decades since mowing was adopted |  |
|  |  | The additional number of different management practices applied at the vineyard level from the 1980s to the 2020s, as a *proxy* of the trajectory toward more integrated weed management (*e.g.* if tillage was the only management practice used in the 1980s and if mowing and tillage were used in the 2020s, the value would be ‘1’ additional weed management practice applied) |  |
| 2020s weed management | | The categorical variable of 4 groups of vineyards sharing the same 2020s weed management:  **‘Chem’:** vineyards with chemical weeding of both rows and inter-rows (n=4)  **‘Till.IR.Chem.R’:** vineyards with tilled inter-rows and chemically weeded rows (n=19)  **‘Mow.IR.Till.R’:** vineyards with mowed inter-rows and tilled rows (n=11)  **‘Till’:** vineyards with tilled rows and inter-rows (n=6) |  |
| Functional differences between the 1980s and the 2020s weed communities | The shift in Community Weighted Means of traits, phenological characteristics and Ellenberg indices | **Traits:**  Specific leaf area (SLA), seed mass, maximum height, leaf Dry Matter Content (LDMC), lateral spread  % of C4  **Phenological characteristics:**  Flowering onset, duration of flowering,  **Ellenberg indices:**  Temperature, light, nitrogen and soil moisture | $\Delta CWM=\frac{{CWM}_{2020's}-{CWM}_{1980's}}{{CWM}_{2020's}+{CWM}_{1980's}}$ |
|  | The shift in CSR strategies | C, S and R scores | $\Delta Score=\frac{{Score}_{2020's}-{Score}_{1980's}}{{Score}_{2020's}+{Score}_{1980's}}$ |
|  | Shift along the functional space | **First dimension is driven by phenology (Dim1)**  From earlier and longer flowering to later and shorter flowering communities | $\Delta CWM1{=Dim1}_{2020's}-{Dim1}_{1980's}$  With Dim1, the coordinates of communities along the first axis of the CWM PCA |
|  |  | **Second dimension driven by soil moisture (EIV_F) and temperature requirements (EIV_T) (Dim2)**  From low soil moisture and high-temperature requirements to high soil moisture and low-temperature requirements | ${\Delta CWM2=Dim2}_{2020's}-{Dim2}_{1980's}$  With Dim2, the coordinates of communities along the second axis of the CWM PCA |
|  |  | **Third dimension is driven by photosynthetic pathways (Dim3)**  From communities dominated by C3 species to communities dominates by C4 species | ${\Delta CWM3=Dim3}_{2020's}-{Dim3}_{1980's}$  With Dim3, the coordinates of communities along the third axis of the CWM PCA |
|  |  | **Fourth dimension is driven by SLA (Dim4)** | ${\Delta CWM4=Dim4}_{2020's}-{Dim4}_{1980's}$  With Dim4, the coordinates of communities along the fourth axis of the CWM PCA |
| Taxonomic differences between the 1980s and the 2020s | Abundance, richness, Shannon and Pielou indices |  | $\Delta Indice=\frac{{Indice}_{2020's}-{Indice}_{1980's}}{{Indice}_{2020's}+{Indice}_{1980's}}$ |

| Table S2 Community Weighted Means contribution to the first four dimensions of the functional PCA of 1980s and 2020s communities (%). SLA, Specific Leaf Area ; LDMC, Leaf Dry Matter Content ; Lat_spr, Lateral spread ; EIV_F, Ellenberg indice for soil moisture ; EIV_T, Ellenberg indice for temperature ; EIV_N, Ellenberg indice for nitrogen ; EIV_L, Ellenberg indice for light; Flow_on, flowering onset ; Flow_dur, duration of flowering. | | | | |
| --- | --- | --- | --- | --- |
|  | Dim1 (%) | Dim2 (%) | Dim3 (%) | Dim4 (%) |
| SLA | 7 | 0 | 36 | 4 |
| LA | 5 | 4 | 3 | 11 |
| LDMC | 7 | 0 | 10 | 10 |
| Seed_mass | 2 | 8 | 10 | 2 |
| Max_height | 9 | 6 | 7 | 3 |
| C4 | 3 | 2 | 2 | 41 |
| Lat_spr | 2 | 17 | 3 | 2 |
| EIV_F | 1 | 21 | 6 | 14 |
| EIV_T | 8 | 20 | 1 | 5 |
| EIV_N | 7 | 4 | 17 | 1 |
| EIV_L | 7 | 13 | 3 | 5 |
| Flow_on | 19 | 5 | 0 | 2 |
| Flow_dur | 20 | 0 | 0 | 0 |

| Table S3 Pairwise correlations between functional metrics at the community levels. SLA, Specific Leaf Area ; LDMC, Leaf Dry Matter Content ; Lat_spr, Lateral spread ; EIV_F, Ellenberg indice for soil moisture ; EIV_T, Ellenberg indice for temperature ; EIV_N, Ellenberg indice for nitrogen ; EIV_L, Ellenberg indice for light ; Flow_on, flowering onset ; Flow_dur, duration of flowering ; LA, leaf area. | | | | | | | | | | | | |
| --- | --- | --- | --- | --- | --- | --- | --- | --- | --- | --- | --- | --- |
|  | LDMC | Seed_mass | EIV_F | EIV_T | EIV_N | EIV_L | Flow_on | Flow_dur | C4 | Lat_spr | Max_height | LA |
| SLA | 0,09 | -0,15 | 0,06 | **-0,18** | **0,35** | -0,09 | **-0,25** | **0,42** | 0,00 | **-0,21** | **-0,20** | **-0,38** |
| LDMC | - | **-0,19** | **0,21** | -0,11 | 0,03 | -0,07 | **-0,34** | **0,22** | **-0,20** | -0,07 | **-0,33** | **-0,31** |
| Seed_mass |  | - | -0,11 | 0,03 | **-0,23** | 0,00 | **0,39** | **-0,34** | **0,21** | **0,26** | **0,38** | **0,32** |
| EIV_F |  |  | - | **-0,48** | **0,21** | **-0,30** | 0,02 | 0,13 | -0,01 | **0,18** | -0,08 | -0,05 |
| EIV_T |  |  |  | - | **-0,30** | **0,64** | **0,20** | **-0,45** | **0,29** | -0,02 | 0,00 | 0,14 |
| EIV_N |  |  |  |  | - | -0,14 | **-0,34** | **0,27** | **-0,45** | -0,13 | -0,07 | **-0,24** |
| EIV_L |  |  |  |  |  | - | **0,33** | **-0,43** | **0,21** | -0,06 | **0,27** | 0,15 |
| Flow_on |  |  |  |  |  |  | - | **-0,72** | **0,56** | **0,34** | **0,75** | **0,42** |
| Flow_dur |  |  |  |  |  |  |  | - | **-0,27** | **-0,29** | **-0,49** | **-0,40** |
| C4 |  |  |  |  |  |  |  |  | - | **0,23** | **0,34** | **0,24** |
| Lat_spr |  |  |  |  |  |  |  |  |  | - | **0,23** | **0,27** |
| Max_height |  |  |  |  |  |  |  |  |  |  | - | **0,51** |

The correlation coefficients of significant correlations are reported in bold. *p*-values were corrected from multiple testing using Holm correction.

| Table S4 Selected fixed effects of the Community Weighted Mean relative differences in the 1980s and the 2020s (SLA, Specific Leaf Area). | | | |
| --- | --- | --- | --- |
|  | **ΔSLA** | **Δ Max_Height** | **Δ Lateral_spread** |
| *Selected predictors* | *No fixed effects selected* | *No fixed effects selected* | *No fixed effects selected* |
| Observations | 108 | 108 | 108 |
| Marginal R^2^ / Conditional R^2^ | 0 / 0.52 | 0 / 0.39 | 0 / 0.12 |

| Table S5 Selected fixed effects of the Community Weighted Mean relative differences in the 1980s and the 2020s (EIV_L, Ellenberg indice for light ; Δ.EIV_T, relative changes in temperature needs ; ΔEIV_N, relative changes in nitrogen needs). | | | |
| --- | --- | --- | --- |
|  | **ΔEIV_L** | **ΔEIV_T** | **ΔEIV_N** |
| *Selected predictors* | *No fixed effects selected* | *No fixed effects selected* | *No fixed effects selected* |
| Observations | 108 | 108 | 108 |
| Marginal R^2^ / Conditional R^2^ | 0.000 / 0.31 | 0 / 0.34 | 0 / 0.44 |

| Table S6 Selected fixed effects of the Community Weighted Mean relative differences in the 1980s and the 2020s (ΔEIV_F, relative changes in soil moisture needs). | | | |
| --- | --- | --- | --- |
|  | **ΔEIV_F** | **ΔFlowering onset** | **ΔFlowering duration** |
| *Selected predictors* | *No fixed effects selected* | *No fixed effects selected* | *No fixed effects selected* |
| Observations | 108 | 108 | 108 |
| Marginal R^2^ / Conditional R^2^ | 0 / 0.31 | 0 / 0.34 | 0 / 0.29 |
