## Appendix S3 for "Climate and management changes drove more stress-tolerant and less competitive plant communities in 40 years"

### Appendix 3

### Title

Climate and management changes drove more stress-tolerant and less competitive plant communities in 40 years

**Authors**

Marie-Charlotte Bopp^1*^, Elena Kazakou^1^, Aurélie Metay^2^, Jacques Maillet^3^, Marie-Claude Quidoz^1^, Léa Genty^2,5^, Guillaume Fried^4^

**Ecological monographs**

**Section S1: Farmer interviews to collect management practices**

The first interview was realised in June 2020 on the weed management practices applied from 1978 to June 2021 (45 minutes) and the second one was conducted in June 2021 to collect information about the more recent weed management practices (15 minutes). The questionnaire was composed of 61 questions with open and multiple-choice answers divided into three sections: (i) farm characteristics (area, wine valorisation, labels), (ii) vineyard characteristics (topography, vine density, grape variety, fertilisation, manure, irrigation) and (iii) weed management practices of rows and inter-rows from 1978 to 2021 (chemical weeding, tillage and mowing).

**Section S2: multivariate analyses to describe climate and management change over time**

To describe climate and weed management changes over the four decades, we computed (i) a Principal Component Analysis (PCA) based on climate characteristics per decade from the 1980s to the 2020s (1980s, 1990s, 2000s, 2010s and 2020s) and (ii) a Principal Coordinates Analysis (PCoA) based on weed management characterisation per decade from the 1980s and 2020s that characterised the trajectory of weed management. In addition to defining the weed management trajectory during four decades, 2020s weed management was also characterised.

**Section S3: classification of vineyards according to 2020s weed management**

To describe the current weed management practices, a PCA was performed on the following agricultural practice variables averaged over the 2015-2021 period (Appendix S1: Figure S1): (i) the annual number of chemical weeding, tillage and mowing applied at the row of vines and the inter-row (*i.e.* the free space between the rows of vines) level, (ii) fertilisation input and soil amendment. To group the vineyards according to their 2020s practices, hierarchical clustering was then performed on the principle components resulting in the classification of vineyards into four clusters (Appendix S1: Figure S1). Cluster ‘Chem’ gathers four vineyards with chemical weeding of both rows and inter-rows. Cluster ‘Till.IR.Chem.R’ regroups 19 vineyards with tilled inter-rows and chemically weeded rows. Cluster ‘Mow.IR.Till.R’ gathers 11 vineyards with mowed inter-rows and tilled rows. Cluster ‘Till’ groups six vineyards with tilled rows and inter-rows.
